# Integrated exome–transcriptome analysis of a schizophrenia case with an exceptionally low polygenic risk score

**DOI:** 10.64898/2026.11.09.765277

**Authors:** Nikolay V Kondratyev, Alexander S Artyuhov, Vera A Guven, Tatyana V Lezheiko, Olga I Efimova, Kirill A Ulianov, Nour Ebrahim, Anna Y Morozova, Georgiy P Kostyuk, Ekaterina E Khrameeva, Erdem B Dashinimaev, Vera E Golimbet

## Abstract

Schizophrenia liability reflects both common polygenic variation and rarer variants of larger effect. We investigated a post-mortem schizophrenia case (SZ07) with a schizophrenia polygenic risk score below the first percentile among ancestry-matched cases, asking whether individual-level genomic and transcriptomic analyses could help identify potentially novel schizophrenia-relevant variants. We used exome analysis with multiple variant-prediction tools to generate a panel of variants and corresponding candidate genes potentially implicated in the pathological process in SZ07. The prioritised variants were then assessed using RNA sequencing of three cortical regions. Of these, only an *IZUMO4* splice-acceptor variant showed direct RNA-level support. Beyond variant-specific analyses, we characterised broader transcriptomic abnormalities in the SZ07 brain. Pathway analysis of gene-level expression deviations implicated innate immune processes, calcium signalling, synaptic biology, and axon remodelling. Notably, compared with conventional single-sample biological pathway analysis, the analysis we used showed only a weak association with neuronal specificity, suggesting their reduced sensitivity to the main axis of variation related to cellular composition in bulk brain tissue gene expression data. Although the candidate genes showed no significant pathway-level convergence, the broader transcriptomic profile revealed coherent biological alterations largely compatible with the documented history of cerebrovascular disease in SZ07. These results demonstrate how the integration of exome and transcriptome data at the individual level enables the formulation and constraint of hypotheses regarding patient-specific genetic features and their molecular consequences, while highlighting important limitations related to tissue composition, comorbidity, and the choice of reference cohort.

## Introduction

Schizophrenia has a highly polygenic architecture dominated by numerous common alleles each carrying small individual effects ^1^. At the same time, rarer coding variants are capable of conferring substantially greater individual risk. These variants were identified at the exome-wide significance level in genes such as *SETD1A*, *SP4*, and *AKAP11* ^2–4^. In the original SCHEMA study, protein-truncating variants in the ten genes reaching exome-wide significance were estimated to occur in fewer than 1% of patients. However, an excess burden of such variants in evolutionarily constrained genes persisted even after genes reaching the false discovery rate threshold were excluded, suggesting that additional rare-variant risk genes remained to be discovered ^2^. This conjecture is supported by the substantially expanded SCHEMA 2.0 analysis, which identified 16 genes at exome-wide significance and additional genes at the 5% false discovery rate threshold ^4^.

Typically, exome or genome sequencing finds many rare variants in a patient. As testing expands to panels and exomes, interpretation of the discovered individual variants becomes the main and mostly nontrivial challenge. A systematic approach is expected to link the gene-disease relationship and find evidence supporting it, which is often problematic ^5^. The scale of the problem is evident in clinical sequencing cohorts: genome sequencing of 1,007 heterogeneous cases produced phenotype-related variants of uncertain significance (VUS) in 24%, alongside a 21% diagnostic yield ^6^, while 23% of affected individuals in a neurodevelopmental-disorder genome cohort carried VUS despite extensive family-based analysis ^7^.

In the best case scenario, direct functional evidence can help resolve variants that remain clinically uncharacterised. For example, a validated *BRCA2* homology-directed repair assay enabled reclassification of 86% of clinically observed VUSs in the assayed DNA-binding domain ^8^. Similarly, saturation-scale functional measurements of *MSH2* supported reclassification of 74% of evaluated VUSs and were concordant with relevant tumour and clinical features ^9^. However, these successes may not be easy to generalise: such assays often target genes for which disease-relevant molecular functions are well defined and experimentally tractable. No comparable clear functional mechanisms have been established for schizophrenia.

In general, interpretation of variants with predicted functional effects could be assisted by RNA expression data. Population transcriptomic studies established that rare variants can be linked to large expression effects through expression outliers and allele-specific expression ^10^. Rare conserved variants near transcription start sites are enriched among expression outliers and allele-specific measurements narrowed rare regulatory effects to small sets of candidate variants ^11^. These observations support a general strategy in which aberrant expression, splicing, or allelic balance provides functional evidence that a candidate rare variant perturbs gene regulation.

Clinical studies have shown that this strategy can substantially improve Mendelian disease diagnosis. For example, in fibroblasts from 197 individuals with inconclusive exome results, systematic analysis of expression, splicing, and allele-specific expression produced diagnoses in 32 cases; causative findings included noncoding variants and deep-intronic variants missed by exome sequencing ^12^. In another study blood RNA analysis found abnormal splicing for 85 of 257 clinically selected variants, while also showing that some canonical splice-site variants had no relevant effect because the affected exons were normally skipped ^13^. RNA analysis is especially informative when anchored to a candidate RNA-affected variant: hypothesis-driven RNA sequencing provided diagnostic or variant-resolution utility in 69% of probands with candidate DNA findings (45% in whom RNA-seq provided a molecular diagnosis), whereas analysis without a candidate produced substantially fewer findings ^14^. Similarly, RNA-seq resolved 26% of 411 hereditary-cancer VUSs predicted to affect splicing ^15^. Thus, transcriptomics is generally most efficient as a complement to genome-scale DNA testing, although expression outlier analysis can also reveal previously overlooked variants.

No experimental system fully recapitulates schizophrenia, but human brain tissue provides the closest approximation to its native cellular and anatomical context. Post-mortem transcriptomic studies have converged on altered neuronal and synaptic programmes, with higher-resolution analyses localising these changes to specific cortical layers, neuronal subtypes, and coordinated neuron–glia programmes, and increasingly linking them to common and rare genetic risk ^16–19^. These studies revealed substantial regional and molecular heterogeneity while also highlighting the need to distinguish disease-related changes from medication, ageing, and other post-mortem confounders ^20,21^.

Polygenic risk scores (PRSs) summarize common-variant liability and may help identify affected individuals whose disease risk is less readily explained by common variation ^22,23^. Psychiatric PRSs robustly capture group-level differences ^1^. Thus, under a liability-threshold model, cases with low PRS could be enriched for high-effect rare variants. Supporting this idea, schizophrenia cases carrying risk-associated copy-number variants (CNVs) have shown lower PRS than noncarriers, particularly for higher-risk CNVs ^24,25^, though the evidence for the association is not universal ^26^. Low PRS therefore provides a rationale for investigating other genetic contributors, rather than a reliable indicator of high-effect rare variation.

Our primary aim in the present study was to determine whether rare, potentially high-impact genetic variants could be revealed and supported by detectable molecular consequences in matched brain tissue. We therefore investigated a schizophrenia donor with an exceptionally low SZ-PRS and available post-mortem brain tissue, integrating exome sequencing with matched transcriptomic analyses to evaluate candidate rare variants and their molecular consequences.

## Results

### Identification of a schizophrenia case with an exceptionally low SZ-PRS in the post-mortem brain collection

During routine genotyping of donors in the post-mortem brain collection (n = 18), we examined schizophrenia polygenic risk scores ^1^. SZ07 was identified as a case of an individual with unusually low SZ-PRS. His standardised score was evaluated against the independent RMHRC case distribution (1221 patients/762 controls), in which it falls below the first percentile (z = −2.03 relative to controls, z_case_=-2.67), i.e. 0.25% (fitted-normal percentile, 0.38%) of cases have a PRS this low or lower, indicating a low common-variant liability to schizophrenia for SZ07, discordant with manifest illness (Figure 1A). Combining the fitted case and control PRS distributions with an assumed lifetime schizophrenia prevalence of 0.7% yielded a PRS-based risk estimate of 0.15%, approximately 79% below population prevalence (bootstrap 95% CI for the relative reduction, 70–85%). The rarity of the score alone, however, is not what motivated individual-level investigation. This case was pursued because of the conjunction of an extreme low polygenic score with available matched post-mortem brain tissue, a combination substantially rarer than either feature alone.

**Figure 1.**
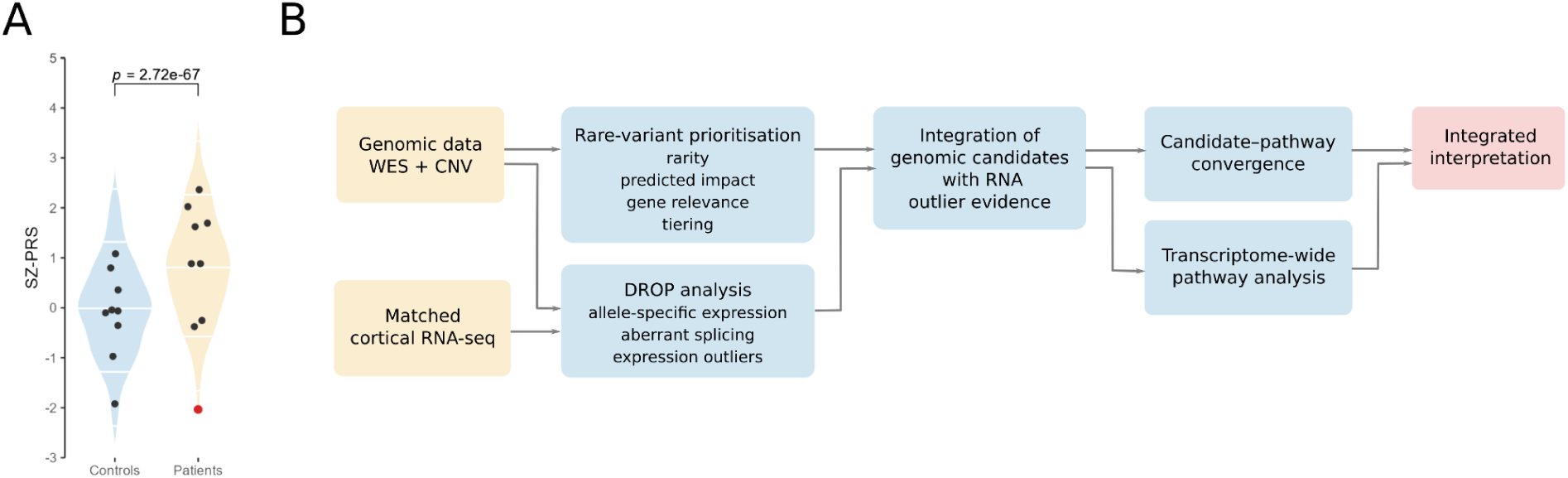
Polygenic-risk context and integrated genomic–transcriptomic analysis of SZ07. **(A)** Distribution of SZ-PRS values in patients with schizophrenia and healthy controls. Violin plots show schizophrenia polygenic scores calculated from the PGC3 results for schizophrenia with the LDpred-2 tool for unrelated and ancestry-matched individuals in the RMHRC sample: schizophrenia patients and healthy controls. SZ-PRS values were standardised relative to controls. White lines indicate medians, deciles, and percentiles of the distributions. Black dots represent scores for individuals from the Skolkovo brain collection, the red dot marks the score of the SZ07 patient. The p-value for the Student’s t-test on the background RMHRC sample is shown. **(B)** Overview of the integrated genomic–transcriptomic analysis. Whole-exome sequencing (WES) and copy-number variant (CNV) data were used for rare-variant prioritisation based on allele frequency, predicted functional impact, gene-level relevance and tiering. Matched cortical RNA-seq was analysed using the DROP workflow to identify allele-specific expression, aberrant splicing and expression outliers, with genomic variant data additionally informing allele-specific analyses. Prioritised genomic candidates were then integrated with RNA outlier evidence. Downstream analyses independently assessed candidate–pathway convergence and transcriptome-wide pathway alterations, which were subsequently considered together in the integrated interpretation.

This patient presented with persistent positive symptoms such as hallucinations and delusions, particularly of a paranoid nature, corresponding to a typical presentation of the disease (ICD10 F20.01). The condition manifested when the patient was in his early twenties. He did not have treatment-resistant schizophrenia; his psychiatric condition was managed with usual doses of antipsychotics, showing partial improvement but with challenges in social and cognitive domains. However, he was socially active until his hospitalisation in his late thirties, had a family, finished his college-level education and was able to work for more than a decade since then. Apart from psychosis, the patient’s medical history was notable for chronic obstructive bronchitis and, later in life, substantial cardiovascular and cerebrovascular disease, including chronic cerebrovascular ischaemic disease with vascular encephalopathy and cognitive impairment, together with cerebral/systemic atherosclerosis and hypertension. No other unusual comorbidities or laboratory findings were documented. Taken together, the disease profile of the patient falls well within the typical range for schizophrenia, providing no clinical indication of an atypical form that might account for the discordantly low polygenic burden. We have matching independent clinical records and DNA samples of SZ07 in the RMHRC and the Alexeev hospital, so the diagnosis was supported by independent clinical records and structured assessment.

This discordance between manifest illness and minimal common-variant burden motivated the working hypothesis that the phenotype might reflect rare, high-impact genetic variation rather than aggregated polygenic liability, prompting exome-level interrogation of the same individual as a single case. We therefore examined whether SZ07’s genome harboured such variation, while recognising that a low score does not require such a variant to exist. An overview of the integrated genomic-transcriptomic analysis is shown in Figure 1B.

### Prioritisation of rare genomic variants in SZ07

First, we examined the patient’s genome for copy-number abnormalities by the array comparative genomic hybridisation. The only interval meeting the predefined aCGH aberration criteria was found in a CNV-dense 6p22 region, containing genes *HFE*, *TRIM38*, and multiple histone-coding genes, none of which were described in the literature as a schizophrenia risk factor (Supplementary Figure 1). CNV scan with the PennCNV tool based on raw genotyping-array intensities did not reproduce this finding. PennCNV also did not reveal any unique CNVs except the gene-poor duplication region of the hg19 chr9:24414621-24505147, the region was not implicated in schizophrenia.

We then examined whether the patient’s exome harboured potentially impactful variants that could be linked to schizophrenia. Whole-exome sequencing data were analysed to prioritise rare coding and splice-region variants, which were stratified into three tiers according to their predicted functional impact, yielding a small set of high-priority candidates (Table 1, Supplementary Table 1). The prioritised variants were classified into three tiers: Tier A included variants with high-confidence LOFTEE loss-of-function annotations or predicted splice-disrupting effects according to SpliceAI (yielding 19 variants); Tier B included damaging missense variants in constrained genes predicted by AlphaMissense and/or REVEL (2 variants); and Tier C included rare protein-altering variants in curated disease-relevant gene sets based on SCHEMA, ASC, and DDG2P (9 variants). Among these, the AKAP11 p.(Tyr1367Cys) was particularly notable as a potentially damaging missense variant in a gene supported by rare-variant associations with both schizophrenia and bipolar disorder ^2,27^, including FDR-significant evidence in the expanded SCHEMA 2.0 schizophrenia analysis ^4^. Other notable rare and potentially impactful variants included LIMK1 p.(Ile32Met), HEY1 c.249+15A>C, and CACNA1D p.(Cys1824Arg). The corresponding genes, *LIMK1* ^28^, *HEY1* ^29,30^, and *CACNA1D* ^31,32^, have established roles in neuronal or neurodevelopmental processes that may be relevant to the pathophysiology of psychosis. However, no clearly high-impact variants in established schizophrenia genes were identified, warranting further evaluation of the functional significance of the prioritised variants.

**Table 1.** Rare protein-altering and predicted splice-altering variants.

| Gene | Variant | Consequence | AF | LoF | SpliceAI | AM | REVEL | LOEUF | SCHEMA | DDG2P |
|---|---|---|---|---|---|---|---|---|---|---|
| **Tier A** |  |  |  |  |  |  |  |  |  |  |
| *ALOXE3* | c.1786-16A>G | Splice PPT intron variant | 6.8e-7 | – | 0.23 | – | – | 1.04 | − | − |
| *ARID3A* | c.*59A>G | 3' UTR variant | 8.2e-5 | – | 0.20 | – | – | 0.44 | − | − |
| *ASCC3* | p.(Ser1563=) | Synonymous variant | – | – | 0.28 | – | – | 0.68 | − | − |
| *CDIN1* | c.273+1G>A | Splice donor variant | 9.9e-6 | HC | – | – | – | 0.93 | − | − |
| *CUZD1* | p.(Ser264Ter) | Stop gained | 1.4e-6 | HC | 0.12 | – | – | 1.07 | − | − |
| *EVC2* | c.3558-2A>G | Splice acceptor variant | 2.1e-6 | HC | 0.92 | – | – | 1.09 | − | + |
| *HEY1* | c.249+15A>C | Intron variant | 2.8e-6 | – | 0.90 | – | – | 0.62 | − | − |
| *ITGA10* | p.(Arg189Ter) | Stop gained | 1.9e-5 | HC | 0.06 | – | – | 0.83 | − | − |
| *IZUMO4* | c.218-2A>C | Splice acceptor variant | 1.7e-5 | HC | 0.70 | – | – | 1.65 | − | − |
| *MAVS* | c.626-94G>C | Intron variant | – | – | 0.20 | – | – | 0.90 | − | − |
| *MMS19* | p.(Ala193Gly) | Missense variant | 6.1e-5 | – | 0.39 | 0.31 | 0.39 | 0.79 | − | − |
| *MVD* | p.(Arg228Gln) | Missense variant | 9.2e-6 | – | 0.40 | 0.49 | 0.49 | 1.71 | − | − |
| *MYO5C* | p.(Arg941Ter) | Stop gained | 1.8e-5 | HC | 0.93 | – | – | 0.89 | − | − |
| *NCOA2* | c.3325-181C>T | Intron variant | – | – | 0.21 | – | – | 0.21 | − | − |
| *OLFML2B* | p.(Asn207Ser) | Missense variant | 7.5e-6 | – | 0.45 | 0.08 | 0.05 | 0.94 | − | − |
| *SLC11A1* | c.1315-30C>G | Intron variant | 9.6e-6 | – | 0.30 | – | – | 1.12 | − | − |
| *SPART* | p.(Asp281Glu) | Missense variant | 7.5e-6 | – | 0.27 | 0.19 | 0.26 | 0.76 | − | − |
| *ULK4* | c.1656+1657A>G | Intron variant | 2.3e-4 | – | 0.42 | – | – | 0.94 | − | − |
| *WDR83* | c.-37+187C>T | Intron variant | – | – | 0.24 | – | – | 1.28 | − | − |
| **Tier B** |  |  |  |  |  |  |  |  |  |  |
| *CYTH2* | p.(Arg262Gln) | Missense variant | 1.4e-5 | – | 0.00 | 0.58 | 0.35 | 0.60 | − | − |
| *LIMK1* | p.(Ile32Met) | Missense variant | 7.1e-7 | – | 0.00 | 0.45 | 0.78 | 0.34 | − | − |
| **Tier C** |  |  |  |  |  |  |  |  |  |  |
| *AKAP11* | p.(Tyr1367Cys) | Missense variant | 7.5e-6 | – | 0.00 | 0.56 | 0.50 | 0.32 | + | − |
| *CACNA1D* | p.(Cys1824Arg) | Missense variant | – | – | 0.00 | 0.15 | 0.25 | 0.25 | − | + |
| *DOCK7* | p.(Pro2085Leu) | Missense variant | 5.9e-5 | – | 0.00 | 0.13 | – | 0.55 | − | + |
| *GATA6* | p.(Pro91Ser) | Missense variant | 2.5e-4 | – | 0.00 | 0.06 | 0.33 | 0.53 | − | + |
| *HECTD4* | p.(Val534Met) | Missense variant | – | – | 0.01 | – | – | 0.29 | − | + |
| *KCNQ2* | p.(Pro772Leu) | Missense variant | 1.5e-5 | – | 0.00 | 0.07 | 0.04 | 0.29 | − | + |
| *SLC5A7* | p.(Glu315Ala) | Missense variant | – | – | 0.00 | 0.17 | 0.44 | 0.46 | − | + |
| *SOX10* | p.(Ile203Val) | Missense variant | – | – | 0.00 | 0.08 | 0.27 | 0.25 | − | + |
| *TBX15* | p.(Thr581Ala) | Missense variant | 6.8e-7 | – | 0.00 | 0.05 | 0.20 | 0.47 | − | + |

Variants are grouped according to the prioritisation criteria into Tier A, Tier B, and Tier C. AF, gnomAD v4 allele frequency; LoF, LOFTEE loss-of-function annotation, with HC indicating high-confidence loss of function; SpliceAI, maximum predicted splice-altering score in any of AG/AL/DG/DL scores; AM, AlphaMissense pathogenicity score; REVEL, Rare Exome Variant Ensemble Learner score; LOEUF, loss-of-function observed/expected upper bound fraction. SCHEMA and DDG2P indicate whether the corresponding gene is included (+) (at FDR ≤0.25 for SCHEMA, confidence ≥2 for DDG2P) or not included (−) in the respective curated gene sets; PPT, polypyrimidine tract; UTR, untranslated region. The more detailed report can be found in Supplementary Table 1.

### Functional interrogation of prioritised variants using matched brain RNA-seq

We collected cortical tissue samples from three Brodmann areas (BA4, BA9, and BA22p). RNA-seq data were obtained for 42 cortical samples from 16 donors from the Skolkovo collection: SZ07, seven other donors with schizophrenia, and eight controls (Supplementary Table 2). Most of the genes with prioritised variants had at least 20 counts per sample and a meaningful expression above 10 TPM (Supplementary Figure 2) allowing the potential effects of the variants to be evaluated in the following transcriptome analysis. PCA showed that samples clustered primarily by donor and secondarily by cortical region, with no clear separation by schizophrenia diagnosis (Supplementary Figure 3), consistent with substantial inter-individual variation in human cortical transcriptomes and with the relatively subtle contribution of schizophrenia diagnosis to bulk post-mortem expression variation reported in larger cohorts ^20,33^. Given the individual-level aim and the small cohort with repeated regional sampling, we prioritised outlier analyses against defined reference cohorts. Matched brain RNA-seq and exome data were processed through the Detection of RNA Outliers Pipeline (DROP) framework ^34^ designed to detect monoallelic/allele-specific expression (module MAE), aberrant-splicing (FRASER2), and gene-level expression outliers (OUTRIDER) against the defined cohort background.

The MAE module found only four variants with excess expression of alternate allele in four genes (Supplementary Table 3). Visual inspection in Integrative Genomics Viewer (IGV) showed that three of the four apparent events were artefactual, resulting from incorrect allele assignment. The remaining *IZUMO4* signal was consistent with allele-specific intron retention associated with the prioritised splice-acceptor variant. Genes that contain the reference-skewed variants are even more prone to false positives ^35^. For example, a very strong and consistent effect found in highly relevant *NRGN* gene upon inspection was based entirely on a mapping artefact. The FRASER2 module did not reveal any significant aberrant splicing events, however the *IZUMO4* variant was detected as the highest-ranking non-significant result. Inspection of the highest-ranked nonsignificant splicing events did not reveal additional obvious schizophrenia-related candidates (Supplementary Table 4).

The OUTRIDER module allows identification of individual genes whose expression differs from a supplied cohort. Expression-outlier analyses were restricted to BA9, the region with the largest number of represented donors and an anatomically matched GTEx reference. For this analysis we tested our BA9 group of samples from 15 donors, including two samples from SZ07. Looking into the raw results table it is clear that most consistently changing genes with the largest effects are not arbitrary. Among 48 genes meeting the exploratory magnitude, expression and concordance criteria, 44 showed reduced expression and four showed increased expression in both SZ07 BA9 samples (Supplementary Table 5); none passed OUTRIDER multiple-testing correction. Cell-type specificity based on the Siletti et al. (2023) human brain single-cell atlas ^36^ indicated that 33 of the 48 genes were predominantly associated with the vascular supercluster, including blood-barrier transporters *ABCB1* and *ABCG2*. *FABP3* and *RTN4R* were the only neuron-enriched genes among the 48 selected genes, and both showed increased expression in the two SZ07 BA9 samples.

We processed the local RNA-seq data to maximise compatibility with the GTEx gene-count data and analysed all 19 local BA9 samples jointly with publicly available 269 GTEx BA9 samples ^37^ with OUTRIDER. This approach follows the DROP protocol’s recommendation to supplement small local cohorts with external reference data. In this analysis OUTRIDER revealed *PPP3CA*, a calcium-dependent regulator of synaptic function, as the sole significant expression outlier detected across the two SZ07 BA9 samples, and showed reduced expression in one sample (log₂ fold change = −0.41; adjusted p-value = 0.0063). Notably, *PPP3CA* was among the 16 genes associated with schizophrenia at exome-wide significance in the rare coding variant analysis by Sealock et al. (2026). However, no prioritised coding or splice-altering variant was identified in *PPP3CA* in the SZ07 exome data.

The only prioritised variant with direct support from brain RNA-seq was the rare *IZUMO4* splice-acceptor variant (hg38 chr19:2097250 A>C). The *IZUMO4* splice-acceptor variant produces allele-specific intron retention, yielding a clearly detectable transcript that is predicted to escape NMD (Supplementary Figure 4A) and encoding a predicted altered protein product with a novel 44-residue C-terminus. Although IZUMO4 belongs to the predominantly testis-associated IZUMO protein family, IZUMO4 is the only member of the family that is soluble, potentially secreted, and expressed outside of testis, suggesting a potential neofunctionalisation ^38^. The single-nucleus atlas of Siletti et al. (2023) indicates that IZUMO4 is weakly expressed in most cell types and regions of the brain, but peaks in cortical astrocytes and two specific neuronal subtypes: glutamatergic hippocampal/entorhinal neurons and histaminergic hypothalamic neurons of the tuberomammillary nucleus (Supplementary Figure 4B). But as far as we know a specific brain function for IZUMO4 has not been established, and no convincing evidence currently links it to a psychiatric phenotype. Exploratory analyses of the altered C-terminal sequence identified predicted donor-specific HLA-I-binding peptides; however, composition-matched null analyses showed that neither the predicted binder yield nor microbial sequence similarities exceeded chance expectation (Supplementary Table 6; further details provided in the Supplementary Note). Thus, the variant has a demonstrable RNA-level consequence, but the available data provide no evidence linking it to the schizophrenia phenotype.

### Assessment of cell-type composition in the SZ07 brain

Because the OUTRIDER results could reflect differences in cellular composition, we assessed cell-type marker expression (Supplementary Table 7) and performed CIBERSORTx deconvolution using the neocortex subset of the Siletti et al. (2023) scRNA-seq data. Both approaches indicated reduced neuronal representation in SZ07 than in most other donors across the sampled cortical regions (Figure 2 A, B; Supplementary Table 8). When looking into more detailed cell type representations we did not observe reduced representation of endothelial markers or inferred vascular proportion (Supplementary Figure 5 A, B; Supplementary Table 9). However, both analyses revealed an increase in fibroblast signatures relative to other samples, with consistently elevated fibrotic-ECM marker-based signatures (Supplementary Figure 5 C), while the endothelial core signal is comparatively preserved. This pattern is consistent with chronic vascular/perivascular remodelling rather than a simple loss of endothelial cells.

**Figure 2.**
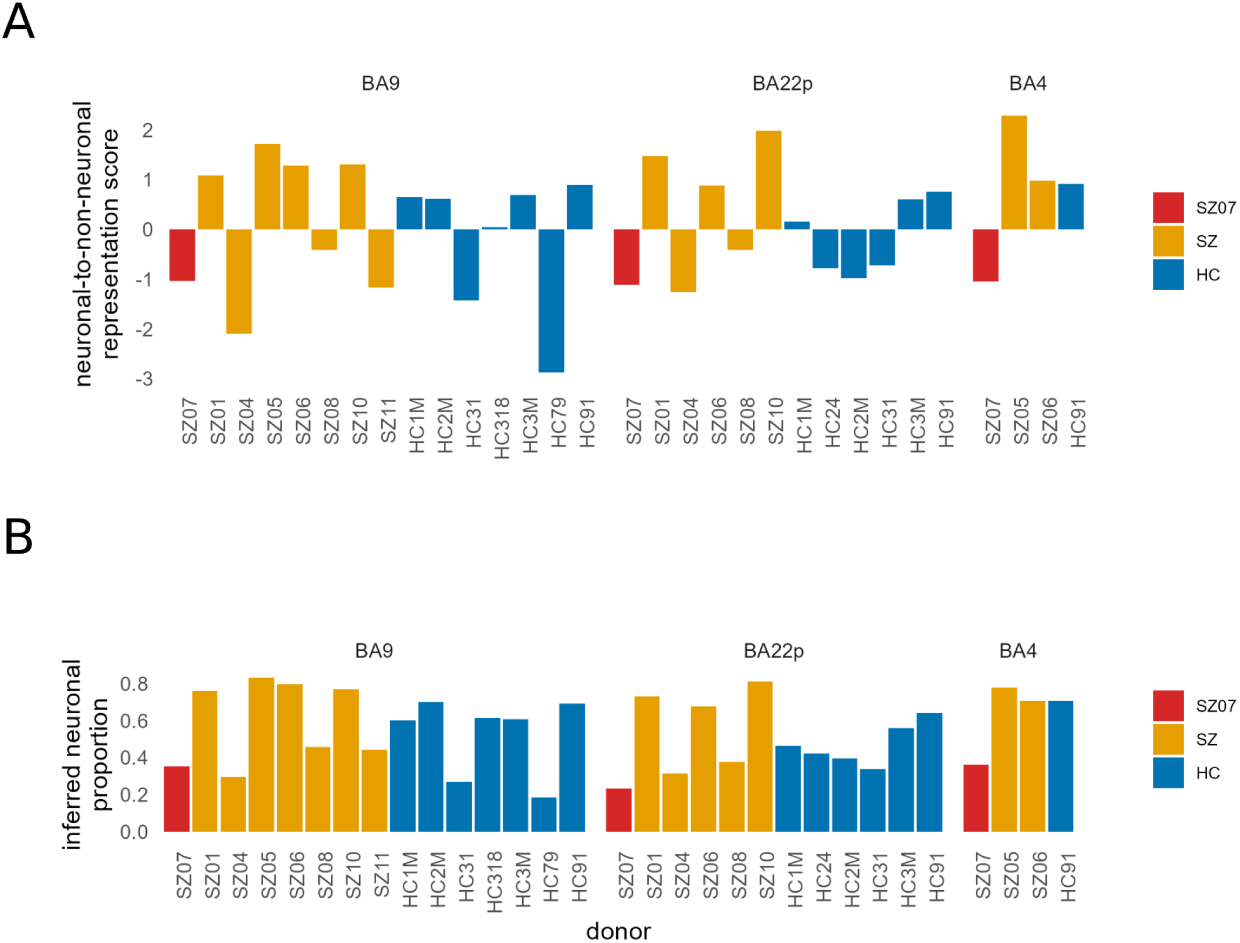
Reduced neuronal representation in the SZ07 cortex. **(A)** Relative neuronal representation, defined as the difference between mean neuronal and non-neuronal marker z-scores, across BA9, BA22p, and BA4. **(B)** Neuronal proportions independently inferred by CIBERSORTx using the cortical subset of the Siletti et al. (2023) reference. In A-B, SZ07 is red, other schizophrenia donors orange, and controls blue.

### Evaluation of pathway-level convergence with prioritised variants

We applied gene set enrichment analysis (GSEA) to OUTRIDER-derived z-scores to identify coordinated expression deviations in SZ07 relative to the local BA9 cohort. By accounting for shared expression patterns, OUTRIDER highlights gene-level deviations that are not readily explained by broader transcriptomic variation, potentially aiding the interpretation of individual rare variants.

Initial assessment based on the in-house data DROP analysis revealed many enriched pathways for SZ07. The strongest result was Hallmark’s “genes regulated by NF-kB in response to TNF” pathway (adjusted p-value = 1.35e-08, NES = 2.3). Other prominent positively enriched clusters include synaptic and neurotransmitter signalling, calcium-ion regulation, toll-like receptor (TLR) & cytokine signalling, humoral/complement immunity, and neuronal-development pathways, whereas negatively enriched gene sets include mitochondrial respiratory-chain and RNA-processing-related processes (Supplementary Figure 6, significant non-redundant pathways are listed in the Supplementary Table 10). The analogous analysis with the external GTEx reference did not produce any significant GSEA hits, probably indicating that SZ07-specific variation was not unique enough in comparison with the larger dataset.

Genes harbouring the tiered variants were evaluated for their representation in significantly enriched pathways identified from OUTRIDER-derived z-scores. Particular attention was given to leading-edge membership, as leading-edge genes contribute directly to the enrichment signal. Tiered variant-containing genes occurring in significant pathways were considered to show biological convergence with the transcriptomic findings. Leading-edge membership provides stronger support than pathway membership alone; however, pathway membership also enables the evaluation of genes with negligible or undetectable expression in the available RNA-seq data. This is particularly relevant for post-mortem brain studies in psychiatric disorders, as disease-related molecular events may occur substantially earlier in life and may no longer be detectable at the time of death.

Fifteen genes harbouring tiered variants were members of the enriched pathways, with KEGG’s “MAPK signalling pathway” (*CACNA1D*) having the smallest p-value = 2.8e-06. *DOCK7*, *NCOA2*, and *SLC11A1* appeared in pathway leading edges (Supplementary Table 11). To assess whether any candidate gene was disproportionately associated with strongly enriched pathways, we considered, for each gene, the smallest single-sample GSEA p-value across its pathways and compared it against a size-matched empirical null, evaluated separately for pathway membership and for leading-edge membership. No tier candidate was over-represented in, or drove, strongly enriched pathways after multiple-testing correction; even the most extreme candidate did not approach significance (smallest unadjusted empirical p*-*value = 0.10, *WDR83*, GO:BP “Response to molecule of bacterial origin”; Supplementary Table 12). So, we detected no statistically supported pathway-level convergence for the prioritised candidates under this analysis.

### Testing pathway-level transcriptomic alterations in SZ07

Compared with other samples in the cohort, SZ07 exhibited a distinctive transcriptional profile characterised by enrichment of calcium signalling, innate immune/TLR–NFκB and complement pathways, together with axonal-regeneration and synaptic-plasticity programmes and suppression of RNA-processing/ribosome-biogenesis pathways (Figure 3A, Supplementary Table 13). Positive NES values for neuronal pathways were noteworthy because they occurred despite the reduced neuronal representation in SZ07 inferred earlier.

**Figure 3.**
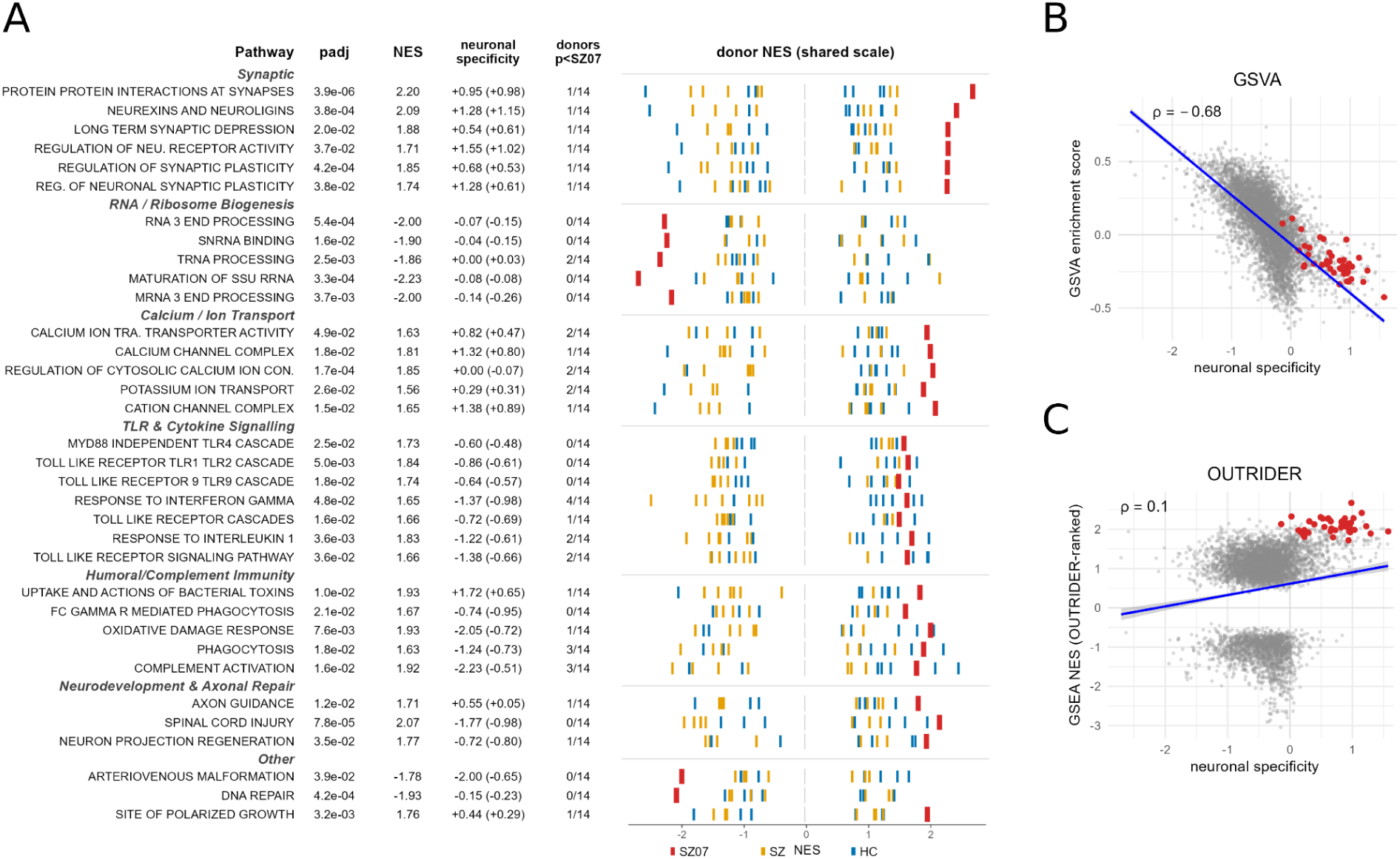
Pathway-level transcriptional changes in the SZ07 brain (A) Donor-wise fgsea results for pathways significant in the OUTRIDER-ranked analysis based on BA9 local data. The most significant results for each functional theme are presented. Columns show adjusted p-values, SZ07 NES, neuronal specificity (mean neuronal-vs-non-neuronal Δlog2 over leading-edge genes, with all pathway genes in parentheses), and the number of other donors with a smaller enrichment p-value than SZ07 (n = 14). The right panel shows donor NES values; pathways are ranked by the NES gap between SZ07 and the next donor. (B) Relationship between pathway “neuronality” measure and SZ07 pathway scores for GSVA across 6,190 pathways. Neuronality is the mean neuronal-vs-non-neuronal Δlog2 of pathway genes. ρ denotes Spearman correlation; lines show ordinary least-squares fits with 95% confidence bands. Red points denote enriched synaptic pathways. (С) Same as in B for the OUTRIDER/fgsea data.

We attempted to reproduce these findings using GSVA, a commonly used single-sample gene-set or pathway-scoring (ssPA) approach (Supplementary Table 14). The effects for the previously highlighted neuronal pathways appeared to be inverted (Supplementary Figure 7). However, SZ07’s conventional GSVA pathway scores were strongly associated with neuronal specificity (Spearman ρ = −0.68, p < 2.2e-16) (Figure 3B), whereas this relationship was much weaker for OUTRIDER-residual-based pathway enrichment (ρ = +0.1, p = 6.21e-14) (Figure 3C). On a common pathway set, an interaction model confirmed that the specificity–score relationship differed between methods (specificity × method β = +1.80, p < 2e-16), reversing the GSVA slope of −1.53 to the weak positive slope (+0.27). The association with neuronal specificity was markedly attenuated in the OUTRIDER-based analysis compared with GSVA, suggesting a potential advantage for investigating individual expression anomalies in bulk brain tissue.

## Discussion

### Polygenic scores as a guide to individual-level investigation

Polygenic scores (PGS), including disease-associated PRS, have research applications beyond clinical screening, for which utility remains context-dependent and routine psychiatric implementation is limited ^39,40^. PGSs can help investigate changes in diagnostic ascertainment ^41^ and the genetic relevance of clinical distinctions, such as mood-incongruent psychosis in bipolar disorder ^42,43^. There are also PGS-informed cellular studies: for example, studies that linked neuronal electrophysiology to diagnosis and donor cognition ^44^, and cross-psychiatric PGS to iPSC morphology and its response to fibronectin ^45^.

Here, unusually low schizophrenia PRS motivated investigation of contributors to disease liability insufficiently captured by common-variant scores. Under an additive liability-threshold model, affected carriers of large-effect rare variants may require less common-variant liability to develop disease. Consistent with this hypothesis, studies across several complex disorders have reported lower polygenic burden among affected carriers of high-effect variants, sometimes with enrichment of monogenic diagnoses among low-PRS cases ^46–51^.

In schizophrenia, evidence comes primarily from CNV studies and varies across study designs. Bergen et al. (2019) reported lower PRS among cases carrying schizophrenia-associated CNVs, particularly higher-risk CNVs ^25^, while Chen et al. (2026) found lower PRS among East Asian cases carrying large deletions ^24^. Taniguchi et al. (2020) observed enrichment of established high-impact variants among CNV carriers in the lowest PRS decile, despite no significant overall PRS difference ^52^. This pattern was less evident in the population-based iPSYCH2015 cohort: Vaez et al. (2026) found neither significantly lower schizophrenia-spectrum PGS among recurrent-CNV carriers nor significant recurrent-CNV–PRS interactions, highlighting ascertainment differences and limited power as possible explanations ^26^. Together, these findings motivate low-PRS case selection as a research strategy, but whether it improves rare-variant discovery or interpretation remains to be established in comparative studies.

### Integrated exome–transcriptome approach to rare-variant prioritisation and functional interpretation

In this study, we employed an integrative, stepwise analytical framework to evaluate potentially pathogenic variation in a single individual. Rare genomic variants were first prioritised using structural-variant and exome-based approaches and were subsequently evaluated using matched brain transcriptomic data. This design allowed candidate variants to be tested with the available transcriptomic context using both an in-house produced cohort and public data with the help of the DROP framework ^34^. As a result, several initially plausible findings were not supported by independent analyses, and the majority of prioritised variants did not show convincing transcriptomic or pathway-level convergence.

One of the most important limitations of this study is the inherent uncertainty of rare-variant prioritisation. Any filtering strategy represents a trade-off between sensitivity and specificity: stringent thresholds reduce the large background of rare variants carried by unaffected individuals, but inevitably increase the risk of excluding genuinely relevant variants ^53^. Accordingly, the tiered strategy used here should be regarded as a means of defining a tractable set of hypotheses rather than identifying all potentially causal variation. Restriction to predicted loss-of-function, splice-disrupting and damaging missense variants may miss context-dependent, gain-of-function, regulatory or other mechanisms not captured by these annotations, while prioritisation based on established disease-gene sets necessarily favours current knowledge and may exclude as-yet undiscovered risk genes.

This is particularly relevant to schizophrenia, for which large-scale exome studies indicate that additional rare-variant risk genes remain to be identified. The original SCHEMA study analysed 24,248 cases and 97,322 controls and identified 10 genes at exome-wide significance ^2^, whereas the expanded SCHEMA 2.0 analysis included 87,959 cases and 150,587 controls but increased this number to only 16 genes ^4^. This illustrates the diminishing returns of sample expansion alone and suggests that many genuine risk genes may remain below achievable statistical thresholds, motivating complementary approaches such as the PRS-based screen idea we used in this study.

This uncertainty provides an important rationale for integrating functional data into variant prioritisation. Computational predictions primarily identify variants with an increased probability of affecting gene or protein function, whereas an observed effect on allelic expression, splicing, or gene expression provides an independent line of evidence that can strengthen or weaken support for a candidate variant ^54^, but their predictions remain affected by gene-, variant- and training-set-specific biases ^55,56^. Moreover, computational approaches are also intrinsically incomplete, as conventional variant-effect predictors perform less well for gain-of-function and other non-loss-of-function mechanisms, which constitute a substantial fraction of dominant Mendelian disease mechanisms ^57,58^. The *IZUMO4* splice variant found in this study provides a potential example: the aberrant transcript is predicted to encode a novel C-terminal sequence and therefore could, in principle, acquire molecular properties not captured by conventional variant-effect prediction, although the downstream immunological mechanism explored here remains unsupported (Supplementary Note).

Reference-panel selection presents an additional practical challenge for RNA-seq-assisted interpretation of rare variants, including VUSs in clinical diagnostics. An individual patient’s expression profile must be evaluated against a background that adequately represents expected variation in the relevant tissue. Small local cohorts may offer closer technical comparability but provide limited power to detect modest expression abnormalities. External datasets can expand the reference population, but differences in tissue composition, physiological state, sample handling and quantification may introduce additional biases. Consequently, increasing reference-panel size does not necessarily improve sensitivity to a particular variant-associated effect. Our analyses illustrate this dependence on the reference cohort. The local and GTEx-augmented analyses produced different expression-outlier and pathway-enrichment results, and the principal pathway findings from the local analysis were not reproduced in the combined cohort. These differences could reflect a broader representation of biological variation, technical heterogeneity, or changes in the variation captured by the latent-factor model. Without independently established positive controls, we cannot determine their relative contributions or identify either analysis as the more accurate representation of variant effects.

For individual variant interpretation, an expression abnormality therefore requires evaluation alongside the predicted molecular consequence, tissue relevance, allelic evidence and clinical phenotype. Conversely, failure to detect an outlier provides limited evidence against pathogenicity unless the assay is sufficiently sensitive to the expected effect.

### Pathway-level expression anomalies and cellular-composition effects

Beyond evaluating individual genes, we used matched brain transcriptomics to investigate coordinated pathway-level expression deviations that might inform the functional interpretation of prioritised variants.

Single-sample gene-set or pathway-scoring (ssPA) methods such as GSVA summarise coordinated expression changes into pathway-level scores and support functional interpretation of individual transcriptomes ^59^. However, their clinical use for variant interpretation remains limited, although related disease-specific transcriptional signatures provide proof of principle. For example, elevated interferon-stimulated gene signatures have supported interpretation of candidate variants in genes associated with type I interferonopathies ^60,61^. Such evidence is likely most informative when the expected downstream phenotype is well established. For example, a conceptually related approach uses ssPA of variant-induced transcriptional profiles in *in vitro* models as a functional readout ^62^. Whether generic single-sample pathway scores can provide reproducible evidence for variant interpretation remains largely unexplored.

A major consideration in single-sample pathway analysis of bulk tissue is that pathway scores can reflect differences in cellular composition rather than altered pathway activity within a given cell type. This limitation is not specific to GSVA or to absolute-expression-based approaches. Indeed, ssGSEA-derived scores have been successfully used to infer stromal, immune, and other cell-type abundances from bulk transcriptomes, demonstrating that rank-based gene-set scores remain sensitive to the representation of cells expressing the corresponding genes ^63,64^. More directly, decomposition of bulk RNA-seq profiles has shown that compositional variation can account for a substantial fraction of expression variability and can materially affect pathway-level enrichment estimates ^65^. The problem is particularly relevant in brain tissue, where variation in cellular composition accounts for a large component of bulk transcriptional variation and can differ considerably even between anatomically closely matched samples ^66^. This is consistent with our observation that inter-individual differences represented the largest source of variation in the bulk transcriptomic data. Although estimated cell-type proportions can be included as covariates to mitigate cellular-composition effects, such adjustment may not fully eliminate composition-related confounding in bulk brain data. In particular, Toker et al. (2023) ^67^ showed that cell-type estimates can incompletely account for compositional variation and cautioned against assuming that covariate adjustment removes this signal when cellular composition is strongly confounded with the condition of interest. For the brain it is additionally complicated by the pronounced intra-cell heterogeneity, since bulk transcriptomic measurements reflect not only cellular abundance but also contributions from axonal, dendritic, or synaptic compartments. In the present study, the strong association between neuronal specificity and conventional GSVA scores is consistent with this known sensitivity to tissue composition.

Pathway enrichment based on OUTRIDER-derived z-scores was only weakly associated with neuronal specificity. Because OUTRIDER models shared gene-expression covariation through an autoencoder ^68^, this attenuation is compatible with reduced sensitivity to composition-related variation. However, OUTRIDER does not explicitly model cell identities or proportions, and a weaker association with neuronal specificity does not establish that compositional effects have been removed. The comparison also involves different gene-level inputs and pathway-scoring procedures, so the difference cannot be attributed solely to the autoencoder. Nevertheless, the results suggest that pathway analysis of OUTRIDER-derived expression deviations may complement conventional pathway scoring when investigating coordinated abnormalities in an individual sample. Attribution of these abnormalities to prioritised variants requires additional evidence, because tissue injury and other physiological processes can also generate coherent pathway-level changes.

### Transcriptomic findings in the context of cerebrovascular disease

The documented history of cerebrovascular disease in SZ07 provides a plausible unifying explanation for many of the molecular phenotypes observed in SZ07’s brain. The reduced inferred neuronal abundance and downregulation of mitochondrial programs are consistent with well-described consequences of chronic cerebral hypoperfusion, including neuronal loss, energy failure, and mitochondrial dysfunction ^69^. Genes with strongly reduced OUTRIDER expression scores were enriched for endothelial markers, including *ABCB1* and *ABCG2*, which encode major BBB efflux transporters. Their coordinated reduction is therefore consistent with impaired BBB endothelial identity or function. This interpretation is supported by evidence that chronic cerebral hypoperfusion disrupts the BBB and neurovascular unit and, experimentally, reduces *ABCB1* expression in brain capillaries ^70,71^. *FABP3* showed concordant, subthreshold positive expression deviations in SZ07. Experimental studies implicating FABP3/H-FABP in ischaemic neuronal injury make this finding compatible with injury-related changes ^72,73^. Cerebral ischaemia and ischaemia-reperfusion can also induce endothelial phenotypic transitions, including endothelial-to-mesenchymal transition, providing one possible mechanism by which reduced endothelial identity could coexist with increased mesenchymal/fibroblast-like signatures ^74^. After cerebral ischaemia, toll-like receptors TLR2 and TLR4 are activated in response to DAMPs, and through intracellular pathways such as TAK1/MAPKs or NFκB/AP-1 drive production of cytokines and chemokines. The IL-1 response set fits the same axis ^75^.

Chronic ischaemia can produce a mixed state in which degenerative synaptic changes and an attempted reparative/plasticity program occur simultaneously. Neuronal-development and axonal-repair pathways (axon guidance, spinal cord injury, neuron projection regeneration) fit the post-ischaemic reparative/remodelling program that constitutes a growth-promoting response ^76–79^. The accompanying enrichment of synaptic, calcium-channel, and transporter-related functions may reflect altered neuronal excitability and post-ischaemic plasticity and are also compatible with compensatory remodelling under chronic ischaemia. Despite reduced inferred neuronal representation, SZ07 showed coordinated positive enrichment of presynaptic vesicle release and recycling, postsynaptic receptor/scaffold organization, NMDA-receptor trafficking, and bidirectional synaptic-plasticity pathways. The leading-edge signal was dominated by core synaptic machinery including *STX1A*, *SNAP25*, *STXBP1*, *SYT1*, *RAB3A* and *DLG4*, rather than by generic neurodevelopmental genes. Because experimental chronic cerebral hypoperfusion generally reduces synaptic vesicle-fusion proteins and impairs presynaptic release, the relative transcriptional enrichment of this machinery in SZ07 may represent a compensatory program in surviving neurons. SZ07 additionally shows NMDA-receptor subunit remodelling involving *GRIN2C* and *GRIN2D*. Of particular relevance, experimental ischaemia selectively increases GluN2C/GRIN2C without comparable increases in GluN2A or GluN2B, and GluN2C-containing receptors can limit calcium influx and reduce vulnerability to global cerebral ischaemia ^80^. Together with concurrent enrichment of excitatory and inhibitory synaptic components, these findings are consistent with homeostatic network remodelling rather than a nonspecific increase in excitatory transmission.

It is tempting to hypothesise that the genetic background reflected in SZ07’s unusually low SZ-PRS might also influence compensatory responses to ischaemic injury. Human genetic studies provide some biological plausibility for this, although they do not establish a relationship between schizophrenia liability and stroke recovery. The GISCOME study identified genetic associations with 3-month post-stroke functional outcome that persisted after adjustment for baseline stroke condition ^81^, whereas the GENISIS study found little detectable overlap between genetic susceptibility to ischaemic stroke and early neurological change during the first 24 hours after stroke ^82^. Common variants were estimated to explain approximately 7–9% of variance in early neurological change in the initial GENISIS analysis. Subsequent GENISIS work implicated neuronal excitability through ADAM23 and GRIA1 and found suggestive enrichment of sodium-ion transmembrane transport, learning/memory, abnormal CNS synaptic transmission, and several cortical tissues ^83^. A longitudinal GWAS of domain-specific recovery in 1,270 patients reported suggestive associations of motor recovery with axon-guidance genes including *ROBO1*, *ROBO2*, and *PTPRS* ^84^, while GISCOME implicated vascular and recovery-related genes such as *TEK* and *NTN4*. Thus, the molecular phenotype observed in SZ07 converges on biological systems independently implicated in human genetic studies of post-stroke outcome, including synaptic excitability, axonal remodelling, neural plasticity, and vascular responses. This convergence supports the interpretation of SZ07’s synaptic and axonal-remodelling enrichment as reflecting, at least in part, an active response to ischaemic injury rather than neuronal composition alone. Available post-stroke outcome GWASs are currently (2026) too small to estimate genetic correlation with schizophrenia liability with useful precision; consequently, whether the genetic determinants of schizophrenia liability and ischaemic recovery are correlated, and in what direction, remains unresolved.

### Conclusions

Several limitations constrain the interpretation of this analysis. First, prioritization of rare variants remains inherently incomplete, as current prediction methods have limited sensitivity, and the absence of whole-genome sequencing further restricts the detection of potentially significant regulatory or other noncoding variants. Second, although post-mortem brain samples represent the closest and most accessible approximation to native disease-relevant tissue, they reflect a late and substantially distorted state. This state is largely shaped by ageing, treatment, agonal processes, and comorbid pathology rather than by the diagnosis-relevant molecular state at the disease onset. The relatively small local cohort of brain samples further limits the characterisation of normal interindividual transcriptomic variability, since a “one versus reference” design cannot by itself distinguish disease-associated anomalies from individual or incidental molecular features. This problem is particularly relevant to case SZ07, as pronounced cerebrovascular pathology constitutes a powerful competing source of transcriptomic effects that may mask or mimic schizophrenia-related signals. Finally, the motivating premise of the study itself is probabilistic. An extremely low schizophrenia polygenic risk score increases the motivation to search for alternative sources of genetic predisposition, but it does not imply that a large-effect rare variant necessarily exists. Case SZ07 may simply represent an individual who developed schizophrenia despite an unusually low measured burden of common variants. Thus, given these limitations, we can neither exclude a contribution of rare genetic factors nor, with comparable confidence, attribute the phenotype to such a contribution without more compelling evidence.

In conclusion, this study illustrates the importance of carefully evaluating observed molecular anomalies when determining disease causes at the individual level. Although population studies are essential for defining the genetic architecture of schizophrenia, individual-level approaches address how this risk manifests in a particular patient. Integrating knowledge obtained at the population level with genomic analysis and analysis of functional outliers may help reveal patient-specific molecular hypotheses that remain hidden in group comparisons. Despite substantial methodological limitations, such approaches may facilitate the translation of psychiatric genomics from population-level discoveries to clinically meaningful individual assessment.

## Methods

### Clinical data

Diagnoses of schizophrenia were made according to the criteria of the International Classification of Diseases, 10th Revision (ICD-10) adopted in Russia and were confirmed by a structured psychiatric interview: the Mini-International Neuropsychiatric Interview (MINI). The study was approved by the Ethics Committee of the Russian Mental Health Research Center (18 March 2022, No. 873).

### Post-mortem brain samples

Post-mortem human brain samples from patients were collected in the Alexeev Clinic No. 1 and samples from healthy donors were obtained via National BioService Russian Biospecimen CRO (St. Petersburg, Russia). Informed consent for using human tissues for research was obtained in writing from all donors or their next of kin by the tissue provider bank. All control subjects died suddenly with no prolonged agony state and had no history of neurological and psychiatric disorders or alcohol and drug abuse. The brain blocks selected for molecular analysis were reported by the tissue providers as free of gross focal pathology. This assessment did not constitute a systematic neuropathological evaluation of cerebrovascular disease. Post-mortem brain frontal blocks 1-1.5 cm thick were stored at -80°C wrapped in foil and placed in zip-lock bags. For sample dissection, The Atlas of the Human Brain (AHB) ^85^ was used to locate the areas of interest, a posterior part of the superior temporal gyrus of the left hemisphere (posterior Brodmann area 22 (BA22p), Wernicke’s area), the dorsolateral prefrontal cortex (Brodmann area 9, BA9), and the primary motor cortex (Brodmann area 4, BA4). Frontal blocks with surfaces nearest to AHB level: 60 (MNI: -36.57; ICL: 34.82; MCP: 20.08) were placed into the cryostat Leica CM1950 chamber at -20°C before dissection for 15-20 min on custom-made aluminium support. A tissue sample weighing approximately 150 mg was cut out from a selected area using a metal scalpel. Dissected samples were then collected with tweezers, put into tubes, and placed on dry ice immediately. All materials used during dissection (scalpels, tweezers, tubes) were sterile and chilled using dry ice before use.

### SZ-PRS computation

Single nucleotide polymorphism (SNP)-based Polygenic Risk Scores (PRS) were calculated with data obtained from Illumina GSA chip (v2). Genetic data were imputed with help of the Michigan Imputation Server by the Minimac4 tool with the HRC r1.1 (European panel). PRS for schizophrenia ^1^ was inferred with the LDpred2-auto tool ^86^ available in the R package “bigsnpr” (v1.12.18). As a sample for comparison we employed an in-house ancestry-matched RMHRC sample (762 controls, and 1,221 patients, excluding SZ07; all unrelated). We used leave-one-out summary statistics that excluded the subset of Moscow samples enrolled in the PGC3 schizophrenia study ^1^; the original summary statistics were used for all other samples.

Rarity of SZ07’s SZ-PRS among schizophrenia cases calculation (*x*_ind_ is individual PRS, *μ*_case_ is mean SZ-PRS among cases and *σ*_case_ is the standard deviation among cases).

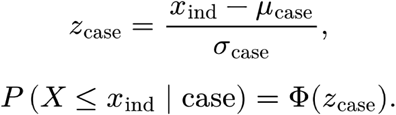

To estimate PRS-associated schizophrenia risk, Gaussian distributions with separately estimated means and standard deviations were fitted to the case and control scores. Distributional fit was assessed using normal Q–Q plots. The fitted probability densities were combined with an assumed lifetime schizophrenia prevalence of K = 0.7% using Bayes’ theorem:

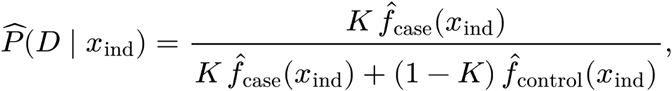

where *f̂*_case_ and *f̂*_control_ are the fitted case and control probability densities. Relative risk reduction versus population prevalence was calculated as

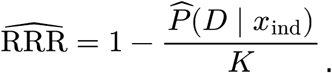

These estimates describe risk associated with the PRS alone, without conditioning on SZ07’s known diagnosis. Percentile bootstrap 95% confidence intervals were obtained from 10,000 resamples drawn separately within the case and control groups, refitting both distributions in each iteration while holding SZ07’s score and *K* fixed.

### CNV analysis

Patient DNA was analysed for genome-wide copy-number abnormalities using SurePrint G3 Human CGH 8×60k Microarray (Agilent Inc., USA) with NA10851 used as a reference DNA. CGH data were analysed with Feature Extraction software, v11.0.1.1 and DNA copy number changes were defined with Agilent Genomic Workbench software using the ADM2 algorithm (threshold = 6.0). Gains and losses were determined involving at least three consecutive probes and mean log2(test/reference) ratio of ≥ 0.25 or ≤ −0.25, respectively.

Copy-number variants (CNVs) were detected from raw Illumina GSA IDAT data from 497 ancestry-matched, unrelated individuals using GenomeStudio and PennCNV. IDAT files were processed in GenomeStudio using the corresponding GSA backbone manifest and cluster files, and per-marker log R ratio and B-allele frequency values were exported. A study-specific population B-allele frequency file was generated from 100 randomly selected samples, and an hg19-based GC-content model was constructed to account for genomic-wave effects. CNVs were called using containerised PennCNV (v1.0.5) (`genomicslab/penncnv`) with the standard `hhall.hmm` model and GC correction ^87^. Downstream analyses were restricted to autosomal CNVs supported by at least 10 SNPs, and CNVs observed in a single individual were prioritised for further interpretation.

### Whole-exome sequencing

Exome sequencing was performed on the patient’s DNA extracted from whole blood (2×150 PE reads, x100 coverage) with SureSelect Human All Exon V8 (Agilent Inc., USA), NextSeq550 (Illumina Inc, US). Rare variants identified by whole-exome sequencing were functionally annotated with Ensembl VEP and prioritised after excluding variants with a gnomAD population maximum allele frequency ≥0.001, and recurrent internal artefact sites. Variants with fewer than four reads supporting the alternate allele, or missing/invalid allelic-depth information, were excluded. Remaining variants were classified into three tiers. Tier A represented predicted loss-of-function or splice-disrupting variants, defined by high-confidence LOFTEE annotations ^88^ or a maximum SpliceAI delta score ≥0.20 ^89^, respectively. Tier B comprised predicted damaging missense variants in genes constrained against predicted loss-of-function variation (gnomAD v4.1 LOEUF < 0.6), defined by an AlphaMissense ^90^ likely-pathogenic classification (score ≥0.564; the threshold is from the original paper) or REVEL ≥0.773 ^91^, using the ClinGen-recommended threshold ^55^. Tier C included rare protein-altering variants in genes belonging to curated disease-relevant gene sets, defined by SCHEMA schizophrenia gene-burden results at FDR ≤0.25 ^2^, Autism Sequencing Consortium results at FDR ≤0.25 ^92^, or DDG2P/Genomics England PanelApp entries with confidence ≥2. BipEx gene-burden statistics ^27^ were retained as annotations but were not used to assign Tier C membership.

### RNA isolation and library preparation

RNA-seq libraries were prepared in two batches, Batch 1 (RMHRC) and Batch 2 (Skolkovo). Total RNA was isolated using RNeasy column kits (Qiagen, USA) in Batch 1, followed by DNase I treatment (NEB, USA) and purification with the CleanRNA Standard kit (Evrogen, Russia), with RNaseOUT ribonuclease inhibitor (Invitrogen, USA) included during isolation to suppress RNase activity; for Batch 2, total RNA was isolated using TRIzol (Invitrogen, USA) according to the manufacturer’s protocol. Qubit fluorometer (Thermo, USA) was used to assess RNA quantity and integrity (Qubit RNA HS & Qubit RNA IQ Assay, Thermo, USA) for Batch 1 and an Agilent 2100 Bioanalyzer (Agilent Technologies, USA) for Batch 2. Both preparations used approximately 200 mg of brain tissue. RNA-seq libraries were prepared using strand-specific, poly(A)-enrichment protocols: Batch 1 libraries were generated from 500 ng of total RNA using the NEBNext Ultra II Directional RNA Library Prep Kit together with the NEBNext Poly(A) mRNA Magnetic Isolation Module (NEB, USA), whereas Batch 2 libraries were generated from 1 μg of total RNA using the TruSeq Stranded mRNA kit (Illumina, USA). Libraries were sequenced in 2 × 150-bp paired-end mode, targeting approximately 20 million individual reads per sample, using an Illumina HiSeq X instrument for Batch 1 and an Illumina HiSeq 4000 instrument for Batch 2. RNA-seq availability varied across donors and cortical regions because RNA extraction was unsuccessful for some tissue samples. Samples showing discordance between RNA-derived genotypes and the corresponding donor DNA genotypes were excluded because their donor–region assignments could not be reliably established.

### RNA-seq data processing

RNA-seq reads were processed using a Snakemake workflow designed to generate analysis-ready data for the Detection of RNA Outliers Pipeline (DROP) ^34^ with parameters matching the Broad Institute GTEx RNA-seq pipeline ^37^. Reads were adapter- and quality-trimmed with Trim Galore (v0.6.2). Reads were aligned using STAR v2.7.11b in two-pass mode to the GTEx-compatible GRCh38 reference assembly lacking ALT, HLA, and decoy contigs, with GENCODE v47 gene annotation. STAR was run using the parameter set employed by the Broad GTEx RNA-seq pipeline. The alignment step generated coordinate-sorted genomic BAM files, transcriptome-coordinate BAM files, gene-level read counts, and splice-junction counts. PCR and optical duplicates were subsequently marked using Picard MarkDuplicates v2.27.5, and the resulting duplicate-marked BAM files were used as the primary inputs for downstream DROP analyses.

### DROP workflow analysis

Aberrant transcriptomic events were subsequently analysed using DROP v1.6.1 ^34^. Aberrant splicing was assessed with FRASER2 using split and non-split read counts extracted from the STAR-aligned BAM files, whereas allele-specific expression was evaluated with the DROP MAE module by integrating RNA-seq alignments with individual whole-exome sequencing variant calls. Local gene-expression counts were generated from aligned BAM files using the DROP workflow, with the same reference genome and GENCODE v47 annotation used for GTEx v11. In the joint OUTRIDER analysis, local BA9 gene-count data and GTEx count matrices were harmonised to a common set and ordering of gene identifiers. OUTRIDER was run within DROP using the autoencoder implementation, with the encoding dimension selected by optimal hard thresholding (`useOHTtoObtainQ = true`). Candidate allele-specific expression events were visually inspected in Integrative Genomics Viewer ^93^ to assess read-level support and exclude mapping or allele-assignment artefacts.

### Cellular composition inference

Cellular composition was assessed using both reference-free marker expression and transcriptomic deconvolution. The marker-based analysis quantified expression of predefined neuronal, astrocytic, oligodendroglial, microglial, mural (pericytes and vascular smooth muscle cells), fibroblasts and endothelial gene panels (Supplementary Table 7) in each sample to assess whether apparent transcriptional differences could reflect variation in tissue composition. For each marker gene, log_2_CPM values were standardised across donors within each brain region, excluding genes detected in fewer than three donors or with no expression variance. Gene-level z-scores were then averaged within each marker panel for each donor. Scores therefore represent relative enrichment or depletion of a cell-type or functional transcriptional signature within a brain region.

For deconvolution, an adult human cortical single-cell reference ^36^ was available for composition estimation with CIBERSORTx deconvolution ^94^. We used the neocortex subset of these data in two deconvolution analyses: (1) neuronal versus non-neuronal fractions and (2) combined neuronal fraction and supercluster-level of other non-neuronal cell-type fractions (“Astrocyte”, “Oligodendrocyte”, “Oligodendrocyte precursor”, “Committed oligodendrocyte precursor”, “Microglia”, “Vascular”, “Fibroblast”, “Ependymal”, “Bergmann glia”, “Choroid plexus”). We used `--single_cell TRUE --rmbatchSmode TRUE --QN FALSE` parameters for the analysis.

Siletti et al. (2023) data were used to construct pathway “neuronality” scores. For each gene we computed a cell-type specificity value as the log₂ fold-difference in expression between neuronal and non-neuronal cells (Δlog₂), derived from the reference data; positive values indicate neuron-enriched genes. Each pathway’s neuronality score was the mean Δlog₂ across its member genes (genes absent from the reference omitted), so that positive scores denote neuron-enriched pathways and negative scores non-neuronal ones.

### Post-OUTRIDER pathway enrichment analysis

Genes were ranked according to their OUTRIDER expression-outlier zScore statistics and analysed by gene set enrichment analysis (GSEA) ^95^. For the initial and prioritised variant-based analysis, the metric for GSEA was constructed from z-scores from individual samples using a conservative signed-minimum statistic with discordance shrinkage:

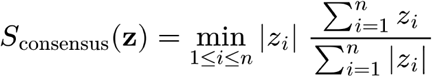

For exploratory analysis of subthreshold expression deviations, we selected genes with concordant log₂ fold-change directions, absolute OUTRIDER z-scores above the sample-specific 90th percentile, and corrected mean expression >30 in each of the SZ07 BA9 samples.

For comparison with data from other donors, use of this metric introduces a bias depending on the number of available samples; therefore, for between-donor comparisons, we used the median z-score instead. For GSEA we used gene-set collections from Hallmark, Reactome, BioCarta, KEGG, Gene Ontology, WikiPathways, and the Human Phenotype Ontology (HPO), as provided through the msigdbr R package (v7.5.1). Pre-ranked gene set enrichment analysis was performed for each donor using the fgsea R package (v1.30.0) ^96^. Donor-specific gene-level statistics were used to rank genes, and gene sets containing 20-200 represented genes were tested for enrichment at either end of the ranked distribution. To reduce redundancy among enriched gene sets, significantly enriched pathways were collapsed using the `collapsePathways` function from the fgsea package. Pathway dependence was assessed using the donor-specific ranked gene statistics, with a dependence threshold of P=0.01, and only the resulting non-redundant main pathways were retained for downstream interpretation.

For candidate-gene prioritisation, pathway and leading-edge membership were first defined from the original GSEA performed on the complete ranked gene list. To prevent circular support arising from a candidate’s own expression statistic, GSEA was then repeated separately for each candidate after removing that gene from both the ranked list and the corresponding pathway gene sets. Candidate–pathway relationships, including leading-edge membership, were retained from the original analysis, whereas pathway enrichment used to evaluate functional convergence was taken from the leave-one-gene-out analysis.

Candidate significance was assessed against an empirical null distribution of exome-testable genes matched for pathway-annotation profile. For each gene, this profile was defined by the number of associated pathways falling within predefined effective pathway-size classes, thereby accounting jointly for pathway annotation burden and the distribution of specific versus broad pathways. Two definitions of candidate–pathway association were evaluated independently. Under set membership, a candidate was associated with all tested pathways containing that gene; under leading-edge membership, it was associated only with significantly enriched pathways in which the gene occurred in the original GSEA leading edge. For each candidate, the minimum leave-one-gene-out enrichment p-value across the relevant pathways was used as the test statistic and compared with the corresponding minimum-p distribution among background genes matched on pathway-size profile. Separate null distributions and matching strata were constructed for the set-membership and leading-edge analyses. Empirical p-values were calculated with a +1 correction as

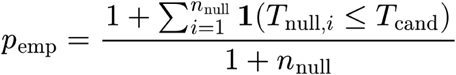

and were Benjamini–Hochberg adjusted separately within the two test classes.

A subsequent donor-specific analysis compared enriched pathways between the index donor and other donors, prioritising pathways with the largest differences in normalised enrichment score (NES). Whole-pathway neuronal specificity was annotated to assess the association of pathway enrichment with neuronal specificity.

To corroborate the OUTRIDER-based enrichment with a method independent of the outlier model, we scored pathway activity with gene set variation analysis (GSVA, v1.52.3) ^97^. Raw counts were filtered to those exceeding 10 counts in at least half the samples, TMM-normalised and log₂-CPM transformed. Genes were mapped to HGNC symbols using the GTF annotation; unmapped genes were dropped and multiple Ensembl IDs mapping to one symbol were summed. Analysis was performed per brain region; biological replicate punches of the same donor and region were averaged on the log-expression scale. Batch was removed with `limma::removeBatchEffect`, preserving diagnosis, sex and age; regions without batch structure were left uncorrected. GSVA scores were computed (Gaussian kernel, gene-set size 20–200) against the same MSigDB collection used for the fgsea analysis. For each pathway, SZ07’s aberrance was quantified as its rank and robust z-score (median/MAD) relative to the other donors, oriented by SZ07’s own direction.

### HLA variants and HLA-binding analysis

HLA class I genotypes were inferred from whole-exome sequencing data using OptiType v1.3.5 ^98^. Trimmed paired-end reads were first enriched for HLA-derived sequences by alignment to the OptiType HLA reference, after which OptiType used integer linear programming to determine the allele combination best supported by the sequencing data, yielding two-field-resolution genotypes for *HLA-A, HLA-B,* and *HLA-C*.

RNA-seq based HLA genotyping was performed using arcasHLA v0.6.0 ^99^. Reads originating from the extended HLA region on chromosome 6 were extracted from the duplicate-marked RNA-seq BAM files and genotyped against an IPD-IMGT/HLA reference using Kallisto-based pseudo-alignment. HLA class I alleles were inferred for *HLA-A, HLA-B,* and *HLA-C*, and class II alleles for *HLA-DRB1, HLA-DQA1, HLA-DQB1, HLA-DPA1,* and *HLA-DPB1*.

The predicted novel C-terminal peptide generated by the *IZUMO4* frameshift was evaluated for potential donor-specific HLA presentation. For HLA class I analysis, the novel C-terminal sequence was tiled into 8-11-mer peptides and evaluated against the donor’s HLA-A, HLA-B and HLA-C alleles using MHCflurry ^100^. Predicted strong binders were collected according to MHCflurry percentile rank. For HLA class II analysis, the complete novel C-terminal sequence was analysed with HLA class II molecules using NetMHCIIpan v4.3 (https://services.healthtech.dtu.dk/services/NetMHCIIpan-4.3) ^101^ using donor-specific HLA-DR, HLA-DQ and HLA-DP genotypes; predicted 15-mer ligands with Rank_EL ≤2 were retained, and their predicted 9-mer binding cores were used for subsequent sequence-similarity analysis.

To determine whether the altered IZUMO4 C-terminal sequence contained more predicted HLA class I binders than expected from its amino-acid composition, we generated composition-preserving shuffled sequences of the altered peptide. For each permutation, amino-acid counts were retained while residue order was randomised, and the sequence was tiled into 8–11-mer peptides and evaluated against the same donor-specific HLA class I alleles using the same strong-binding threshold as for the observed sequence. The number of predicted strong binders was used as the test statistic, and an empirical upper-tail probability was calculated as the proportion of shuffled sequences producing at least as many strong binders as the observed peptide. The implemented analysis used 200 sequence permutations.

Candidate HLA-binding peptides were searched using BLASTP ^102^ against the NCBI non-redundant protein database, restricted by default to bacterial and viral proteins. Full-length peptide matches were classified as exact or near matches; near matches permitted up to two non-identical residues provided that all aligned positions were identical or conservative substitutions, no gaps were present, and the canonical class I anchor positions at P2 and PΩ were conserved.

To assess whether sequence matches obtained by BLAST represented more than the expected background for short peptides of the same composition, a separate composition-preserving null model was applied to each predicted strong-binding peptide. Up to 20 unique shuffled versions of each peptide were generated while preserving its exact amino-acid composition, and the observed and shuffled peptides were searched using identical BLASTP parameters and the same taxonomic search space. Only full-length alignments were retained. Matches were classified as exact or near matches; near matches permitted at most two non-identical residues, required all aligned positions to represent either identities or conservative substitutions, excluded gaps, and required conservation of the class-I anchor positions P2 and PΩ. The observed numbers of total and exact matches were compared with the corresponding distributions across shuffled peptides, with empirical upper-tail probabilities defined as the proportion of shuffled sequences producing at least as many matches as the observed peptide. Recurrence of the same taxa among shuffled peptides was additionally examined as evidence that apparent organism-specific matches reflected sequence-composition background rather than specificity of the observed peptide.

### Use of generative AI tools

Generative AI tools were used during development of the bioinformatic analysis code (Claude Opus 4.7 / 4.8 from Anthropic) to assist with drafting and adapting scripts for the exome and transcriptome analyses, building on pre-existing code and analytical workflows developed by the authors. Generative AI was also used during manuscript preparation (ChatGPT from OpenAI) to assist with language editing and improving the clarity of author-written text. All AI-assisted code and text were reviewed and revised by the authors. The authors retained responsibility for the analytical design, parameter selection, validation and interpretation of results, scientific content, and final wording of the manuscript.

## Data availability

The RMHRC genotypes used for assessing general SZ-PRS distribution are available through Psychiatric Genomics Consortium (“mosc2” and “mosc3” datasets) and are subject to the Consortium’s applicable data-access requirements. Polygenic scores were generated using schizophrenia GWAS summary statistics available at https://pgc.unc.edu. Whole-exome sequencing data for SZ07 are available under controlled access through RMHRC. Any potential sharing of these individual-level genetic data would be subject to applicable legal, institutional and consent-related requirements. RNA-seq data used in this study are available in SRA, accession number PRJNA1520964. GTEx V11 brain-cortex expression counts and metadata used as the external OUTRIDER reference were obtained from the GTEx Portal. Adult human brain single-cell transcriptomic data from Siletti et al. (2023) were used for cell-composition and cell-type-specificity analyses. Publicly available reference and annotation resources used for exome analysis included GRCh38 and GATK known-sites resources, gnomAD, ClinVar, Ensembl VEP and associated functional-prediction resources, and gene-level results from SCHEMA, BipEx, ASC and Genomics England PanelApp. Exact versions, source locations and preprocessing procedures are documented in the accompanying analysis repository.

## Code availability

Custom scripts and reproducible workflows used for whole-exome sequencing, RNA-seq processing, DROP analyses, cell-composition analyses and downstream pathway analyses are publicly available at the project GitHub repository https://github.com/nikolayquadrat/lpb-ex-ome-presession. The repository includes workflows for acquisition and preprocessing of external reference resources, WES alignment and joint variant calling, variant annotation and prioritisation, RNA-seq alignment and quality control, and preparation of inputs for DROP and downstream analyses.

## Author Contributions

Conceptualization: N.V.K. Formal analysis: N.V.K. Investigation (RNA extraction and RNA-seq library preparation): A.S.A., K.A.U., N.E., V.A.G., T.V.L., E.B.D. Investigation (tissue processing): O.I.E. Resources (post-mortem brain procurement): E.E.K., A.Y.M., G.P.K. Data curation: N.V.K., T.V.L., V.E.G. Writing - original draft: N.V.K. Writing - review & editing: N.V.K., O.I.E., E.E.K., E.B.D., V.E.G. Supervision: E.E.K., E.B.D., V.E.G. All authors reviewed the manuscript, approved the final version for publication, and agreed to be accountable for all aspects of the work.

## Acknowledgements

The study was supported by RSF (grant 25-15-00443). The funder played no role in study design, data collection, analysis and interpretation of data, or the writing of this manuscript.

## Competing Interests

The authors declare no competing interests.

## Abbreviations

CNV: Copy Number Variation
COPD: Chronic Obstructive Pulmonary Disease
GSEA: Gene Set Enrichment Analysis
IGV: Integrative Genomics Viewer
NMD: Nonsense-Mediated Decay
NES: GSEA’s Normalised Enrichment Score
PGS: Polygenic Score
PPT: Polypyrimidine Tract
PRS: Polygenic Risk Score
RRR: Relative Risk Reduction
SNP: Single Nucleotide Polymorphism
ssPA: Single-Sample Pathway Analysis
SZ-PRS: Schizophrenia Polygenic Risk Score
TPM: Transcripts per Million
UTR: Untranslated Region
VUS: Variant of Uncertain Significance

## Supplementary Materials

### Supplementary Tables

Supplementary Table 1. Prioritised rare variants discovered in the SZ07 exome and annotation metrics.

Supplementary Table 2. RNA-seq samples inventory.

Supplementary Table 3. MAE results.

Supplementary Table 4. Filtered FRASER2 SZ07-related results.

Supplementary Table 5. Replicate-concordant subthreshold OUTRIDER expression deviations in SZ07 BA9 samples.

Supplementary Table 6. Full-length BLASTP matches of the novel IZUMO4 C-terminal sequence class I binders passing the mimicry-screen filter.

Supplementary Table 7. Marker panels and genes used for transcriptomic scoring of major brain cell types and selected cellular states.

Supplementary Table 8. Deconvolution of RNA-seq samples to neuronal and non-neuronal cell populations.

Supplementary Table 9. Deconvolution of RNA-seq samples to neuronal and major non-neuronal cell types.

Supplementary Table 10. Significant non-redundant gene set enrichment analysis (GSEA) results derived from OUTRIDER gene-level z-scores.

Supplementary Table 11. Tiered genes in GSEA on SZ07 BA9 OUTRIDER results.

Supplementary Table 12. Size-matched minimum-enrichment-p test for tier candidate genes.

Supplementary Table 13. Gene set enrichment analysis (GSEA) results derived from OUTRIDER gene-level median z-scores.

Supplementary Table 14. Per-pathway GSVA scores for SZ07 relative to the donor cohort (BA9).

### Supplementary Figures

**Supplementary Figure 1.**
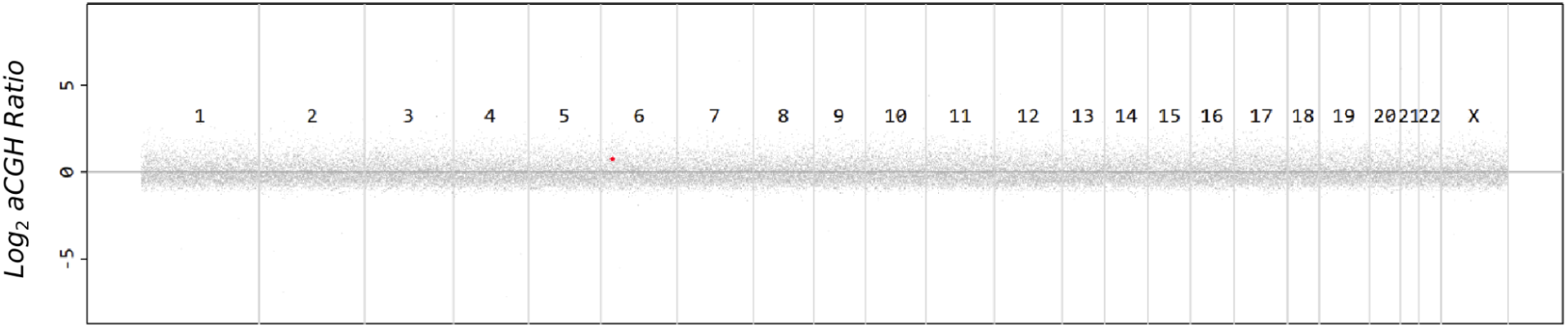
Chromosomal map of aCGH signal for the patient. The aCGH signal was measured as the mean log2 ratio across three probes. The only putative copy-number gain identified by ADM2 signal in the analysis on chromosome six is represented as a red dot.

**Supplementary Figure 2.**
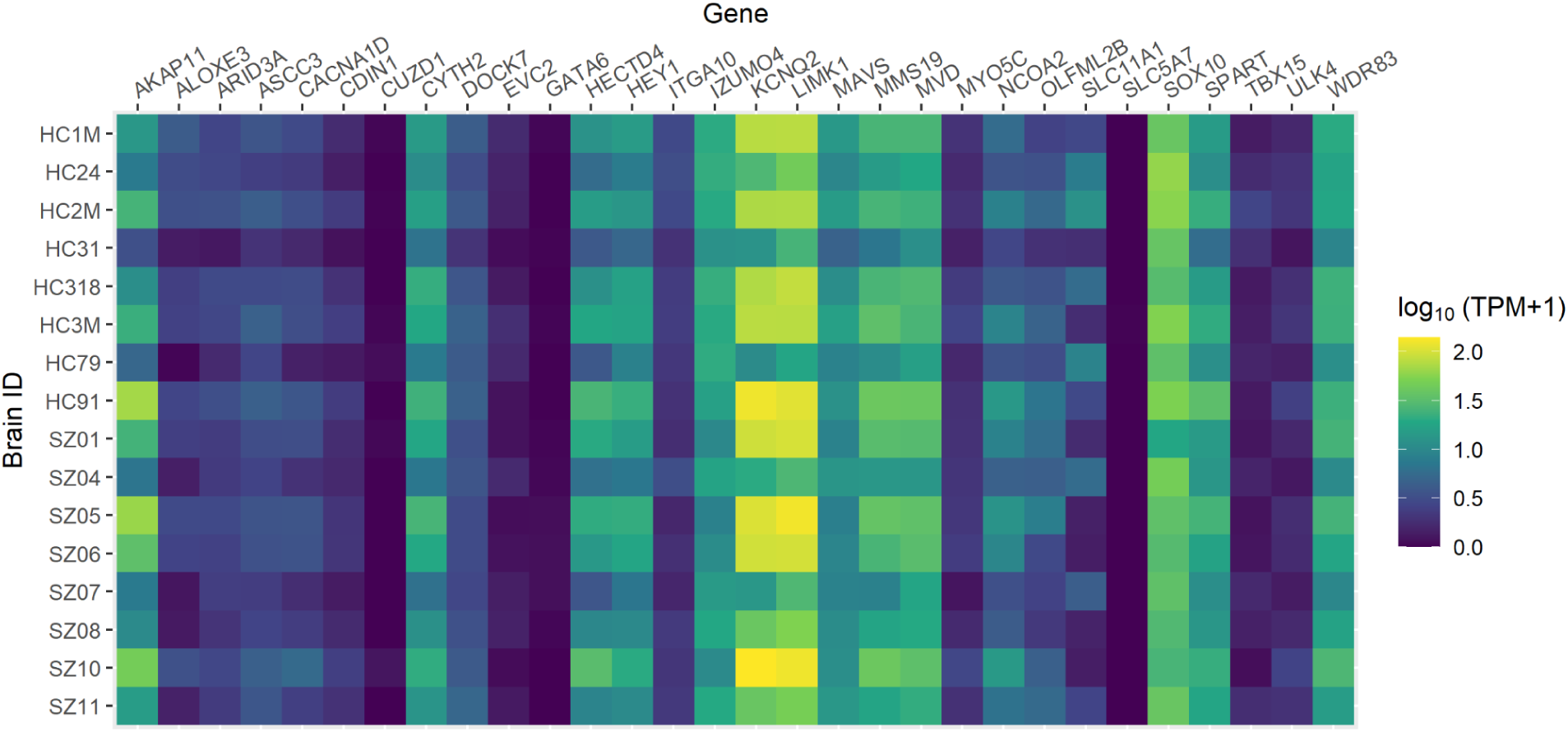
TPM values for genes with the prioritised variants in brain RNA-seq samples. The heatmap shows log_10_ values of the prioritised genes across brain samples, indicating which genes can be directly evaluated using the available RNA-seq data.

**Supplementary Figure 3.**
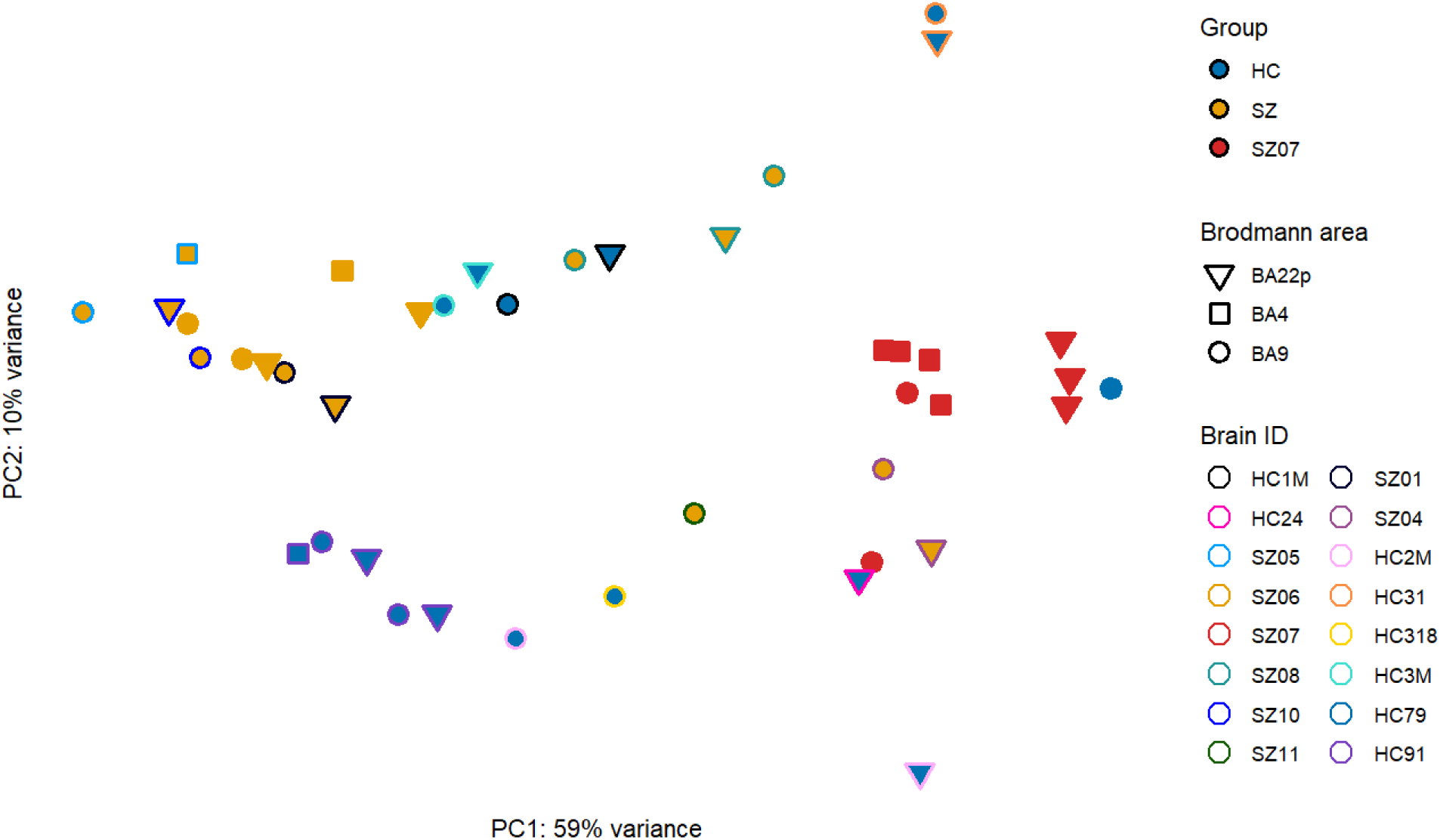
PCA plot of RNA-seq samples used in the current study. PCA was performed on variance-stabilised expression values, with batch and sex effects removed using limma::removeBatchEffect.

**Supplementary Figure 4.**
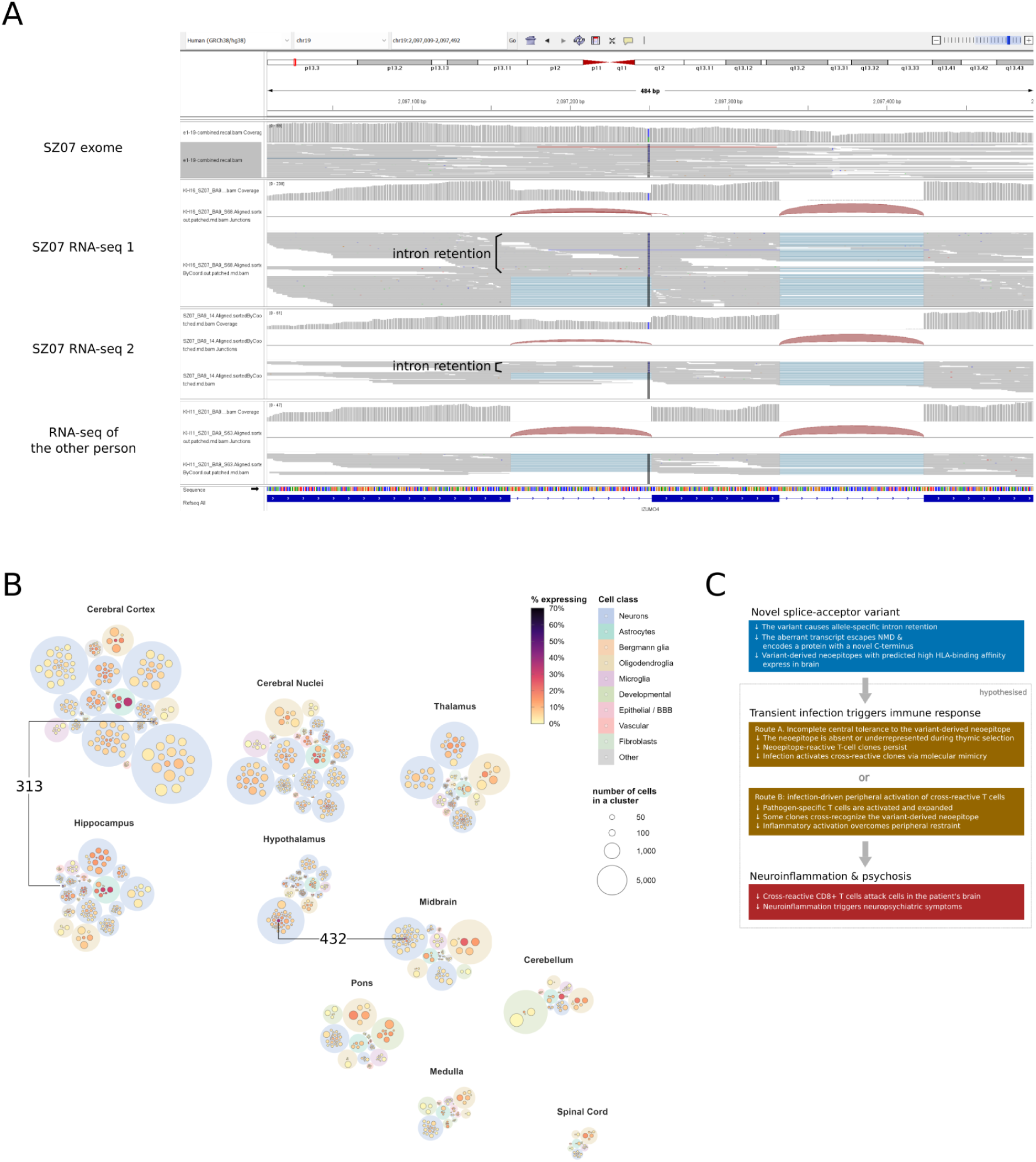
Observed splicing alteration and hypothesised downstream consequences of the IZUMO4 variant. **(A)** IGV visualisation of exome and RNA-seq reads at the *IZUMO4* locus. The vertical line marks the hg38 chr19:2097250 A>C splice-altering SZ07 variant. Both SZ07 RNA-seq replicates show an aberrant splice pattern adjacent to the variant, including intron retention not observed in the control individual. The control RNA-seq sample shows the canonical splice pattern. Red arcs indicate splice junctions, and aligned reads are shown below the coverage tracks. The concordant RNA-seq signal in two SZ07 replicates supports a splice-altering effect of the variant. **(B)** Brain-region distribution and cell-type context of gene expression in the brain single-cell reference from Siletti et al. (2023). Clusters are arranged according to their brain region, with the overall spatial organisation loosely reflecting neuroanatomical relationships. Each circle represents a cluster; circle size indicates the number of cells assigned to that cluster, while circle fill represents the percentage of cells expressing the gene. Clusters belonging to the same supercluster are grouped within a shared background, with background colour indicating broad cell class, including neuronal, glial, vascular, fibroblast, developmental, epithelial/barrier, and other populations. Representative cluster IDs with the highest percentage of cells expressing *IZUMO4* are annotated as in the Supplementary Table 3 from Siletti et al. (2023): 313 – a glutamatergic hippocampal/entorhinal neurons subtype and 432 – histaminergic hypothalamic neurons of tuberomammillary nucleus. Brain-region labels indicate the anatomical origin of each group. **(C)** Proposed mechanism that could potentially link the SZ07 splice variant to neuroinflammation. The splice-altering variant generates an aberrant isoform with a novel C-terminus containing two predicted high-affinity epitopes matched to SZ07 HLA-I alleles, providing a plausible substrate for CD8+ T-cell recognition. Infection could promote pathogenic recognition through incomplete central tolerance to the neoepitopes and/or activation of cross-reactive T-cell clones by molecular mimicry, potentially leading to brain-directed immune responses, neuroinflammation, and neuropsychiatric symptoms.

**Supplementary Figure 5.**
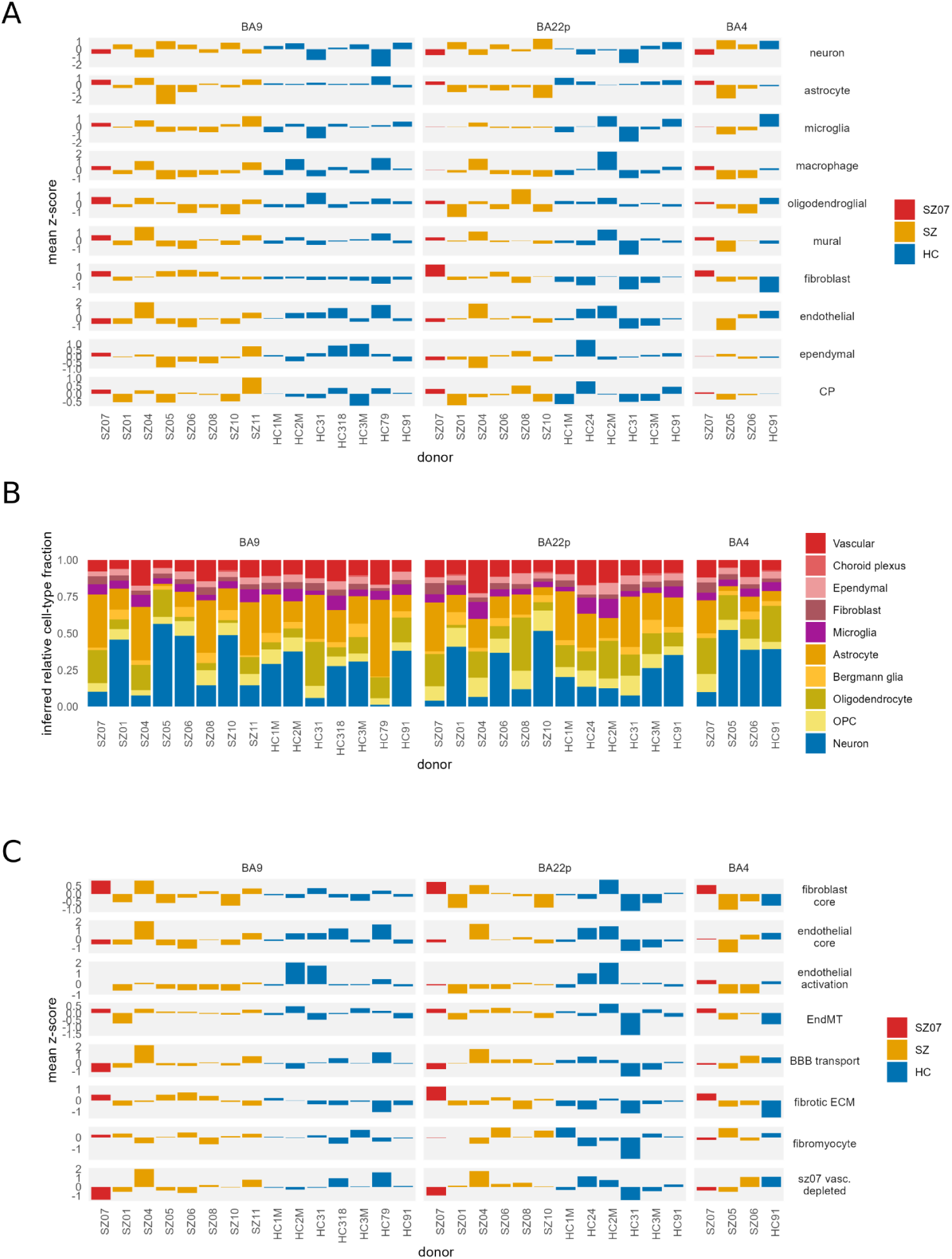
Marker-based and deconvolution-based assessment of cellular composition and vascular signatures across cortical regions. **(A)** Marker-based cell-type representation scores derived from mean gene-level z-scores. SZ07 is highlighted in red, other schizophrenia (SZ) donors in orange, and healthy controls (HC) in blue; **(B)** Relative cell-type fractions inferred by CIBERSORTx using the cortical subset of the Siletti et al. (2023) reference dataset; **(C)** Vascular and perivascular transcriptional signatures across cortical regions. Mean gene-level z-scores are shown for fibroblast core, endothelial core, endothelial activation, endothelial-to-mesenchymal transition (EndMT), blood-brain barrier (BBB) transport, fibrotic extracellular matrix (ECM), fibromyocyte, and SZ07-depleted vascular gene signatures in BA9, BA22p, and BA4. SZ07 is highlighted in red, other schizophrenia (SZ) donors in orange, and healthy controls (HC) in blue. In all panels, bars represent individual donors.

**Supplementary Figure 6.**
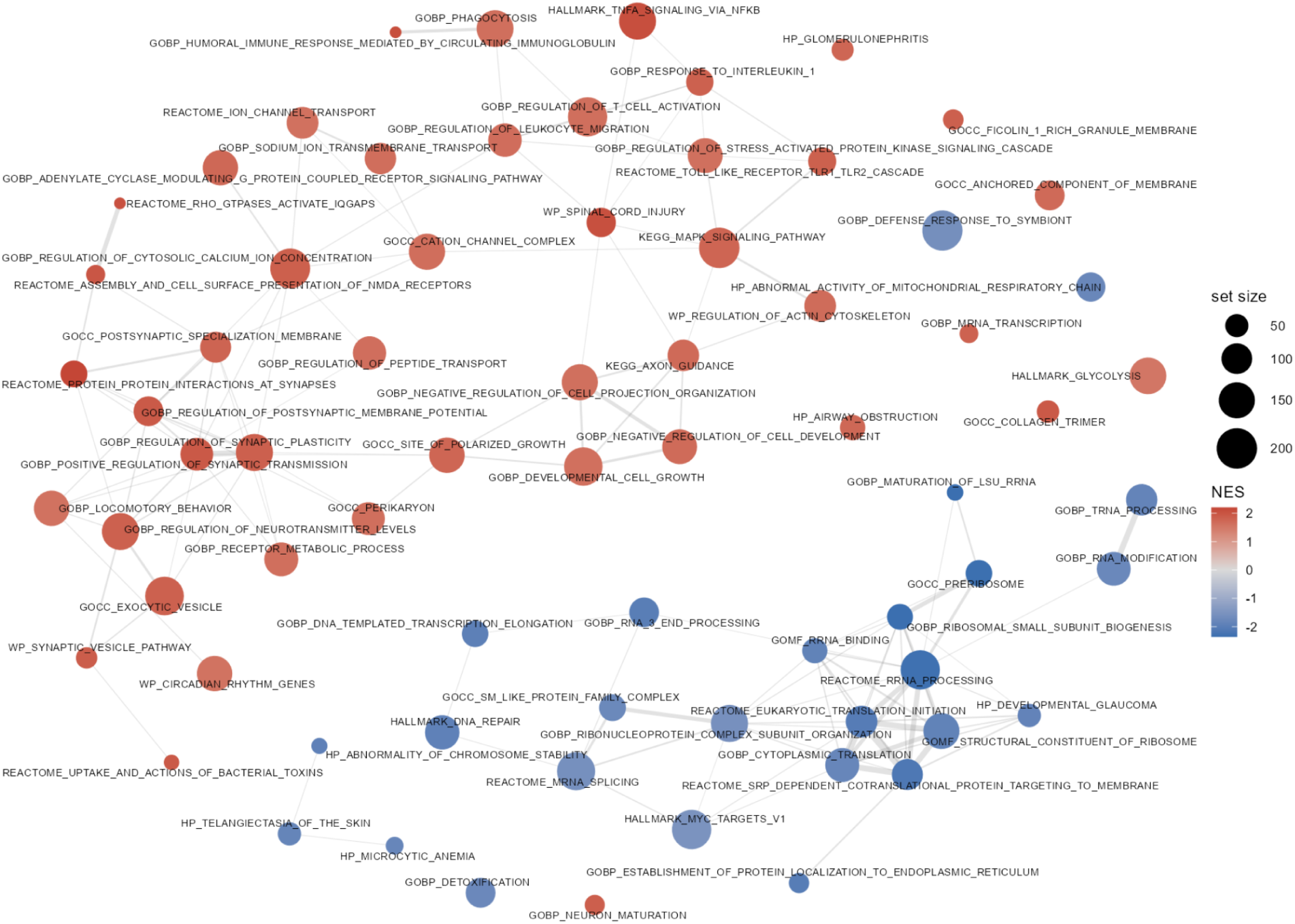
Enrichment map of pathways identified by GSEA of OUTRIDER-derived z-scores. Enrichment map of gene sets associated with aberrant expression in SZ07 identified by GSEA of OUTRIDER-derived z-scores using the local BA9 reference cohort. Node size reflects gene-set size, colour indicates normalized enrichment score (NES; red, positive; blue, negative), and edges indicate gene-set overlap.

**Supplementary Figure 7.**
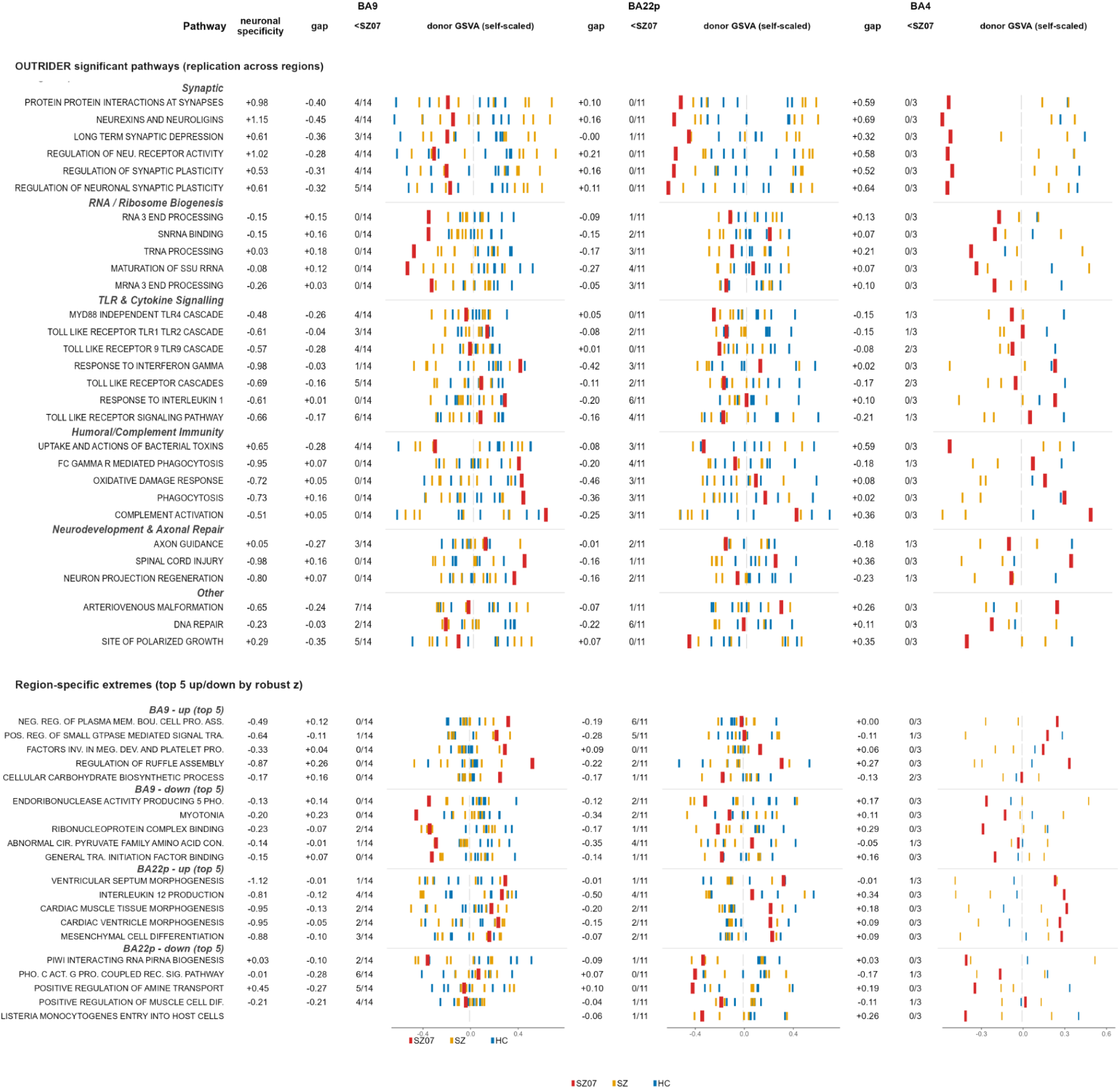
Region-wise GSVA pathway scores for SZ07 relative to the donor cohort. Each row is a pathway; columns show three brain regions (BA9, n = 14 donors; BA22p, n = 11; BA4, n = 3, excluding SZ07). For each region, per-donor GSVA enrichment scores are shown as ticks on a region-specific (self-scaled) axis (SZ07 red, other schizophrenia donors orange, controls blue; vertical grey line at 0), preceded by two summary columns: gap, the difference between SZ07’s score and the most extreme other donor in SZ07’s direction; and <SZ07, the number of other donors whose score is more extreme than SZ07’s in that direction (out of the region’s donor count). The neuronal specificity column (shared across regions) is the pathway’s mean per-gene neuronal-vs-non-neuronal specificity (Δlog₂ from the deconvolution reference; positive = neuron-enriched). The upper block shows the pathways called significant in the OUTRIDER/fgsea analysis, grouped by functional theme; the lower block shows the five most extreme up- and down-scoring pathways (by SZ07 robust z-score, median/MAD relative to other donors) selected separately in BA9 and BA22p. Pathway names are abbreviated.

### Supplementary Note. Potential immunological consequences of the *IZUMO4* splice variant

Because the splice-altering *IZUMO4* variant generated an altered C-terminal sequence, we explored whether the resulting peptides might have unusual donor-specific HLA-binding properties. In silico HLA analysis with HLA alleles inferred from both exome and RNA-seq data (class I: A*11:01/*31:01, B*35:01/*39:01, C*04:01/*12:03) identified two strong MHC class I binders within this novel region for the HLA-A*31:01 allele: RQRDPGAGR (rank = 0.22%, 40 nM) and AQSRLPRQR (rank 0.39%, 94 nM), but no class II binders, motivating a molecular-mimicry/neoantigen hypothesis. Systematic BLAST of the strong binders against the NCBI non-redundant protein database returned several matches in pathogenic bacterial proteins (Supplementary Table 6). Composition-matched null models confirmed that both the strong-binder yield and the exact/near database hits fell within chance expectation. The absence of a post-mortem CD8+/HLA-I effector signature does not exclude a transient immune event around disease onset, which cannot be assessed retrospectively in the available tissue. The neoantigen mechanism was accordingly reframed as a possible transient, onset-era immune trigger that need not persist to post-mortem sampling (Supplementary Figure 4C). SZ07 was diagnosed with COPD, but the available clinical data do not establish whether his condition was associated with a matching opportunistic respiratory pathogen infection (*Stenotrophomonas* or actinomycosis, for example). Thus, although the *IZUMO4* variant has a demonstrable RNA-level consequence, no evidence links it to the schizophrenia phenotype.

The functionally supported *IZUMO4* splice variant raised the possibility of an additional mechanism not captured by conventional variant-effect prediction. The aberrant transcript is predicted to encode a novel C-terminal sequence containing two strong HLA-A*31:01-binding peptides, suggesting that the variant could in principle generate a donor-specific neoepitope. Germline coding variation can generate polymorphic HLA-binding peptides, and altered RNA processing provides an additional source of noncanonical peptide sequences; retained-intron-derived epitopes have been detected directly by immunopeptidomics ^1–4^. The critical unresolved question is therefore not whether such peptides can exist, but whether the *IZUMO4*-derived peptides are naturally processed and presented by the relevant HLA molecule and recognised by a cognate T-cell population.

Such recognition would also depend on cellular and inflammatory context. HLA class I expression in neural cells is normally limited but can be strongly induced by inflammatory signals such as IFNγ, expanding the neuronal immunopeptidome and permitting presentation of otherwise weakly represented antigens. Experimental systems have shown that neuron-specific epitopes can be presented to autologous CD8+ T cells and mediate antigen-dependent neurite injury under inflammatory conditions, while related mechanisms have been demonstrated in oligodendroglial and blood-brain-barrier compartments ^5–7^. Thus, a variant-derived peptide might therefore become immunologically relevant only under particular inflammatory conditions, rather than exerting a constitutive effect.

We further considered whether microbial exposure could provide such a trigger through molecular mimicry. The large diversity of microbial proteomes creates many opportunities for sequence similarity between microbial and self-derived peptides, although sequence similarity alone does not establish functional T-cell cross-reactivity ^8–10^. Infection-triggered cross-reactive immunity is well established in several neurological diseases, and transient infections can precede subsequent brain-directed autoimmune responses. In SZ07, BLAST searches identified several microbial proteins with exact or near matches to the predicted HLA-binding peptides. However, composition-matched null analyses showed that both the number of strong predicted binders and the occurrence of microbial sequence matches were within chance expectation. Thus, the sequence-similarity results alone do not support a specific molecular-mimicry mechanism.

The anatomical expression pattern of *IZUMO4* nevertheless makes the hypothesis biologically interesting. Although expression is generally low in the adult brain, single-nucleus data indicate relatively higher expression in cortical astrocytes and in selected neuronal populations, including histaminergic neurons of the tuberomammillary nucleus. The histaminergic system regulates arousal, sleep-wake state, learning, and memory and has been implicated in neuropsychiatric phenotypes ^11,12^, providing a possible route by which selective immune recognition of *IZUMO4*-expressing cells could have functional consequences. This remains speculative, however, because no brain-specific function of *IZUMO4* has been established and no direct evidence links the gene to schizophrenia.

The available data therefore do not demonstrate that an *IZUMO4*-directed immune mechanism occurred in SZ07. We lack evidence that the predicted peptides were naturally presented, that cross-reactive T cells existed, or that the patient experienced an infection carrying a relevant mimicking antigen. The latter question is particularly difficult to resolve because the available clinical history is incomplete. The absence of a CD8+/HLA-I effector signature in the post-mortem cortex also does not directly test a potentially transient immune event occurring near disease onset. Consequently, the *IZUMO4* finding is best regarded as a biologically plausible but unproven individual-specific mechanism. Given the competing vascular explanation for the post-mortem transcriptomic abnormalities, it cannot presently be assigned a causal role in the psychiatric phenotype.

## Notes

https://github.com/nikolayquadrat/lpb-ex-ome-presession

### Competing Interest Statement

The authors have declared no competing interest.

## References

1. Trubetskoy, V. et al. Mapping genomic loci implicates genes and synaptic biology in schizophrenia. Nature 604, 502–508 (2022).

2. Singh, T. et al. Rare coding variants in ten genes confer substantial risk for schizophrenia. Nature 604, 509–516 (2022).

3. Liu, D. et al. Schizophrenia risk conferred by rare protein-truncating variants is conserved across diverse human populations. Nat. Genet. 55, 369–376 (2023).

4. Sealock, J. M. et al. Analysis of rare coding variation identifies new genetic contributors to schizophrenia. bioRxiv 2026.09.18.752670 (2026) doi:10.64898/2026.09.18.752670.

5. Duzkale, H. et al. A systematic approach to assessing the clinical significance of genetic variants. Clin. Genet. 84, 453–463 (2013).

6. Bertoli-Avella, A. M. et al. Successful application of genome sequencing in a diagnostic setting: 1007 index cases from a clinically heterogeneous cohort. Eur. J. Hum. Genet. 29, 141–153 (2021).

7. Sanchis-Juan, A. et al. Genome sequencing and comprehensive rare-variant analysis of 465 families with neurodevelopmental disorders. Am. J. Hum. Genet. 110, 1343–1355 (2023).

8. Richardson, M. E. et al. Strong functional data for pathogenicity or neutrality classify BRCA2 DNA-binding-domain variants of uncertain significance. Am. J. Hum. Genet. 108, 458–468 (2021).

9. Scott, A. et al. Saturation-scale functional evidence supports clinical variant interpretation in Lynch syndrome. Genome Biol. 23, 266 (2022).

10. Li, X. et al. The impact of rare variation on gene expression across tissues. Nature 550, 239–243 (2017).

11. Montgomery, S. B., Lappalainen, T., Gutierrez-Arcelus, M. & Dermitzakis, E. T. Rare and common regulatory variation in population-scale sequenced human genomes. PLoS Genet. 7, e1002144 (2011).

12. Yépez, V. A. et al. Clinical implementation of RNA sequencing for Mendelian disease diagnostics. Genome Med. 14, 38 (2022).

13. Wai, H. A. et al. Blood RNA analysis can increase clinical diagnostic rate and resolve variants of uncertain significance. Genet. Med. 22, 1005–1014 (2020).

14. Stark, J. C. et al. Clinical applications of and molecular insights from RNA sequencing in a rare disease cohort. Genome Med. 17, 72 (2025).

15. Sha, Y. et al. Refining the interpretation of variants of uncertain significance in hereditary cancer screening through integrated RNA sequencing. Genet. Med. Open 3, 101914 (2025).

16. Fromer, M. et al. Gene expression elucidates functional impact of polygenic risk for schizophrenia. Nat. Neurosci. 19, 1442–1453 (2016).

17. Wu, Y. et al. Developmental and cellular vulnerabilities underlie genetic architecture of schizophrenia. Mol. Psychiatry 10.1038/s41380-026-03862-3 (2026) doi:10.1038/s41380-026-03862-3.

18. Ling, E. et al. A concerted neuron-astrocyte program declines in ageing and schizophrenia. Nature 10.1038/s41586-024-07109-5(2024) doi:10.1038/s41586-024-07109-5.

19. Ruzicka, W. B. et al. Single-cell multi-cohort dissection of the schizophrenia transcriptome. Science 384, eadg5136 (2024).

20. Collado-Torres, L. et al. Regional Heterogeneity in Gene Expression, Regulation, and Coherence in the Frontal Cortex and Hippocampus across Development and Schizophrenia. Neuron 103, 203–216.e8 (2019).

21. Jaffe, A. E. et al. Developmental and genetic regulation of the human cortex transcriptome illuminate schizophrenia pathogenesis. Nat. Neurosci. 21, 1117–1125 (2018).

22. International Schizophrenia Consortium et al. Common polygenic variation contributes to risk of schizophrenia and bipolar disorder. Nature 460, 748–752 (2009).

23. Murray, G. K. et al. Could polygenic risk scores be useful in psychiatry?: A review. JAMA Psychiatry 78, 210–219 (2021).

24. Chen, Y. et al. Contribution of copy number variants to schizophrenia in East Asian populations. Nat. Genet. 1–11 (2026).

25. Bergen, S. E. et al. Joint Contributions of Rare Copy Number Variants and Common SNPs to Risk for Schizophrenia. Am. J. Psychiatry 176, 29–35 (2019).

26. Vaez, M. et al. Recurrent copy number variants and psychiatric outcomes in the context of polygenic scores. JAMA Psychiatry 83, 827–836 (2026).

27. Palmer, D. S. et al. Exome sequencing in bipolar disorder identifies AKAP11 as a risk gene shared with schizophrenia. Nat. Genet. 54, 541–547 (2022).

28. Ben Zablah, Y., Zhang, H., Gugustea, R. & Jia, Z. LIM-Kinases in Synaptic Plasticity, Memory, and Brain Diseases. Cells 10, (2021).

29. Harada, Y. et al. Cell cycle arrest determines adult neural stem cell ontogeny by an embryonic Notch-nonoscillatory Hey1 module. Nat. Commun. 12, 6562 (2021).

30. Than-Trong, E. et al. Neural stem cell quiescence and stemness are molecularly distinct outputs of the Notch3 signalling cascade in the vertebrate adult brain. Development 145, dev161034 (2018).

31. Ortner, N. J., Kaserer, T., Copeland, J. N. & Striessnig, J. De novo CACNA1D Ca2+ channelopathies: clinical phenotypes and molecular mechanism. Pflugers Arch. 472, 755–773 (2020).

32. Shin, J. et al. Cav1.3 calcium channels are full-range linear amplifiers of firing frequencies in lateral DA SN neurons. Sci. Adv. 8, eabm4560 (2022).

33. Johansen, N. et al. Interindividual variation in human cortical cell type abundance and expression. Science 382, eadf2359 (2023).

34. Yépez, V. A. et al. Detection of aberrant gene expression events in RNA sequencing data. Nat. Protoc. 16, 1276–1296 (2021).

35. Stevenson, K. R., Coolon, J. D. & Wittkopp, P. J. Sources of bias in measures of allele-specific expression derived from RNA-sequence data aligned to a single reference genome. BMC Genomics 14, 536 (2013).

36. Siletti, K. et al. Transcriptomic diversity of cell types across the adult human brain. Science 382, eadd7046 (2023).

37. GTEx Consortium. The GTEx Consortium atlas of genetic regulatory effects across human tissues. Science 369, 1318–1330 (2020).

38. Ellerman, D. A. et al. Izumo is part of a multiprotein family whose members form large complexes on mammalian sperm: CHARACTERIZATION OF IZUMO FAMILY PROTEINS. Mol. Reprod. Dev. 76, 1188–1199 (2009).

39. Kullo, I. J. et al. Polygenic scores in biomedical research. Nat. Rev. Genet. 10.1038/s41576-022-00470-z(2022) doi:10.1038/s41576-022-00470-z.

40. Frach, L., David, F. S., Nöthen, M. M. & Forstner, A. J. Polygenic scores in psychiatric research and clinical practice. Med. Genet. 38, 199–210 (2026).

41. LaBianca, S. et al. Changes in genetic contributions to ASD and ADHD by year of diagnosis. JAMA Psychiatry 83, 921–929 (2026).

42. Richards, A. L. et al. Genetic liabilities differentiating bipolar disorder, schizophrenia, and major depressive disorder, and phenotypic heterogeneity in bipolar disorder. JAMA Psychiatry 79, 1032–1039 (2022).

43. Allardyce, J. et al. Association Between Schizophrenia-Related Polygenic Liability and the Occurrence and Level of Mood-Incongruent Psychotic Symptoms in Bipolar Disorder. JAMA Psychiatry 75, 28–35 (2018).

44. Page, S. C. et al. Electrophysiological measures from human iPSC-derived neurons are associated with schizophrenia clinical status and predict individual cognitive performance. Proc. Natl. Acad. Sci. U. S. A. 119, e2109395119 (2022).

45. Coleman, J. R. I. Feasibility and application of polygenic score analysis to the morphology of human-induced pluripotent stem cells. Mol. Genet. Genomics 297, 1111–1122 (2022).

46. Antaki, D. et al. A phenotypic spectrum of autism is attributable to the combined effects of rare variants, polygenic risk and sex. Nat. Genet. 54, 1284–1292 (2022).

47. Huang, Q. Q. et al. Examining the role of common variants in rare neurodevelopmental conditions. Nature 636, 404–411 (2024).

48. Wu, H. et al. Polygenic risk score for low-density lipoprotein cholesterol is associated with risk of ischemic heart disease and enriches for individuals with familial hypercholesterolemia. Circ. Genom. Precis. Med. 14, e003106 (2021).

49. Lu, T., Forgetta, V., Richards, J. B. & Greenwood, C. M. T. Polygenic risk score as a possible tool for identifying familial monogenic causes of complex diseases. Genet. Med. 24, 1545–1555 (2022).

50. Sayadi, A. et al. Rare and common single nucleotide variants in childhood-onset systemic lupus erythematosus. Lupus Sci. Med. 12, e001436 (2025).

51. Baya, N. A. et al. Individuals who deviate from polygenic expectation are enriched for damaging variants in genes linked to rare disease. Am. J. Hum. Genet. 113, 1415–1433 (2026).

52. Taniguchi, S. et al. Polygenic risk scores in schizophrenia with clinically significant copy number variants. Psychiatry Clin. Neurosci. 74, 35–39 (2020).

53. MacArthur, D. G. et al. Guidelines for investigating causality of sequence variants in human disease. Nature 508, 469–476 (2014).

54. Brnich, S. E. et al. Recommendations for application of the functional evidence PS3/BS3 criterion using the ACMG/AMP sequence variant interpretation framework. Genome Med. 12, 3 (2019).

55. Pejaver, V. et al. Calibration of computational tools for missense variant pathogenicity classification and ClinGen recommendations for PP3/BP4 criteria. Am. J. Hum. Genet. 109, 2163–2177 (2022).

56. Rastogi, R. et al. Critical assessment of missense variant effect predictors on disease-relevant variant data. Hum. Genet. 144, 281–293 (2025).

57. Badonyi, M. & Marsh, J. A. Prevalence of loss-of-function, gain-of-function and dominant-negative mechanisms across genetic disease phenotypes. Nat. Commun. 16, 8392 (2025).

58. Gerasimavicius, L., Livesey, B. J. & Marsh, J. A. Loss-of-function, gain-of-function and dominant-negative mutations have profoundly different effects on protein structure. Nat. Commun. 13, 3895 (2022).

59. Toro-Domínguez, D. et al. Benchmarking single-sample gene set scoring methods for application in precision medicine. Brief. Bioinform. 26, bbaf684 (2025).

60. Karla, A. R. et al. SAMHD1 compound heterozygous rare variants associated with moyamoya and mitral valve disease in the absence of other features of Aicardi-Goutières syndrome. Am. J. Med. Genet. A 194, e63486 (2024).

61. Zhang, B. et al. The mutational landscape of STING-induced immunity. Nature 655, 1271–1281 (2026).

62. Thornton, A. M. et al. eVIP2: Expression-based variant impact phenotyping to predict the function of gene variants. PLoS Comput. Biol. 17, e1009132 (2021).

63. Yoshihara, K. et al. Inferring tumour purity and stromal and immune cell admixture from expression data. Nat. Commun. 4, 2612 (2013).

64. Aran, D., Hu, Z. & Butte, A. J. xCell: digitally portraying the tissue cellular heterogeneity landscape. Genome Biol. 18, 220 (2017).

65. Häkkinen, A. et al. PRISM: recovering cell-type-specific expression profiles from individual composite RNA-seq samples. Bioinformatics 37, 2882–2888 (2021).

66. Farahbod, M. & Pavlidis, P. Untangling the effects of cellular composition on coexpression analysis. Genome Res. 30, 849–859 (2020).

67. Toker, L., Nido, G. S. & Tzoulis, C. Not every estimate counts - evaluation of cell composition estimation approaches in brain bulk tissue data. Genome Med. 15, 41 (2023).

68. Brechtmann, F. et al. OUTRIDER: A statistical method for detecting aberrantly expressed genes in RNA sequencing data. Am. J. Hum. Genet. 103, 907–917 (2018).

69. Rajeev, V. et al. Pathophysiology of blood brain barrier dysfunction during chronic cerebral hypoperfusion in vascular cognitive impairment. Theranostics 12, 1639–1658 (2022).

70. Xu, W., Bai, Q., Dong, Q., Guo, M. & Cui, M. Blood-brain barrier dysfunction and the potential mechanisms in chronic cerebral hypoperfusion induced cognitive impairment. Front. Cell. Neurosci. 16, 870674 (2022).

71. ElAli, A., Thériault, P., Préfontaine, P. & Rivest, S. Mild chronic cerebral hypoperfusion induces neurovascular dysfunction, triggering peripheral beta-amyloid brain entry and aggregation. Acta Neuropathol. Commun. 1, 75 (2013).

72. Guo, Q. et al. Fatty acid-binding proteins 3 and 5 are involved in the initiation of mitochondrial damage in ischemic neurons. Redox Biol. 59, 102547 (2023).

73. Wunderlich, M. T. et al. Release of brain-type and heart-type fatty acid-binding proteins in serum after acute ischaemic stroke. J. Neurol. 252, 718–724 (2005).

74. Sun, D. et al. Fluid shear stress induced-endothelial phenotypic transition contributes to cerebral ischemia-reperfusion injury and repair. APL Bioeng. 8, 016110 (2024).

75. Yuan, C., Shentu, Y. & Ji, Q. Research on the innate immune response in transgenic mice following ischemic stroke. Front. Aging Neurosci. 16, 1476913 (2024).

76. Carmichael, S. T. et al. Growth-associated gene expression after stroke: evidence for a growth-promoting region in peri-infarct cortex. Exp. Neurol. 193, 291–311 (2005).

77. Calabresi, P., Centonze, D., Pisani, A., Cupini, L. & Bernardi, G. Synaptic plasticity in the ischaemic brain. Lancet Neurol. 2, 622–629 (2003).

78. Sivilia, S., Giuliani, A., Del Vecchio, G., Giardino, L. & Calzà, L. Age-dependent impairment of hippocampal neurogenesis in chronic cerebral hypoperfusion. Neuropathol. Appl. Neurobiol. 34, 52–61 (2008).

79. Zhang, T. et al. Neuroprotective and axonal outgrowth-promoting effects of tetramethylpyrazine nitrone in chronic cerebral hypoperfusion rats and primary hippocampal neurons exposed to hypoxia. Neuropharmacology 118, 137–147 (2017).

80. Chung, C. et al. Neuroprotection mediated through GluN2C-containing N-methyl-D-aspartate (NMDA) receptors following ischemia. Sci. Rep. 6, 37033 (2016).

81. Söderholm, M. et al. Genome-wide association meta-analysis of functional outcome after ischemic stroke. Neurology 92, e1271–e1283 (2019).

82. Ibanez, L. et al. Overlap in the genetic architecture of stroke risk, early neurological changes, and cardiovascular risk factors. Stroke 50, 1339–1345 (2019).

83. Ibanez, L. et al. Multi-ancestry GWAS reveals excitotoxicity associated with outcome after ischaemic stroke. Brain 145, 2394–2406 (2022).

84. Aldridge, C. M. et al. Genome-Wide Association Studies of 3 distinct recovery phenotypes in mild ischemic stroke. Neurology 102, e208011 (2024).

85. Mai, J. K., Majtanik, M. & Paxinos, G. Atlas of the Human Brain. (Elsevier Science, 2015).

86. Privé, F., Arbel, J. & Vilhjálmsson, B. J. LDpred2: better, faster, stronger. Bioinformatics 10.1093/bioinformatics/btaa1029 (2020) doi:10.1093/bioinformatics/btaa1029.

87. Wang, K. et al. PennCNV: an integrated hidden Markov model designed for high-resolution copy number variation detection in whole-genome SNP genotyping data. Genome Res. 17, 1665–1674 (2007).

88. Karczewski, K. J. et al. The mutational constraint spectrum quantified from variation in 141,456 humans. Nature 581, 434–443 (2020).

89. Jaganathan, K. et al. Predicting splicing from primary sequence with deep learning. Cell 176, 535–548.e24 (2019).

90. Cheng, J. et al. Accurate proteome-wide missense variant effect prediction with AlphaMissense. Science 381, eadg7492 (2023).

91. Ioannidis, N. M. et al. REVEL: An ensemble method for predicting the pathogenicity of rare missense variants. Am. J. Hum. Genet. 99, 877–885 (2016).

92. Satterstrom, F. K. et al. Large-scale exome sequencing study implicates both developmental and functional changes in the neurobiology of autism. Cell 180, 568–584.e23 (2020).

93. Robinson, J. T. et al. Integrative genomics viewer. Nat. Biotechnol. 29, 24–26 (2011).

94. Steen, C. B., Liu, C. L., Alizadeh, A. A. & Newman, A. M. Profiling Cell Type Abundance and Expression in Bulk Tissues with CIBERSORTx. Methods Mol. Biol. 2117, 135–157 (2020).

95. Subramanian, A. et al. Gene set enrichment analysis: a knowledge-based approach for interpreting genome-wide expression profiles. Proc. Natl. Acad. Sci. U. S. A. 102, 15545–15550 (2005).

96. Korotkevich, G. et al. Fast gene set enrichment analysis. bioRxiv (2016) doi:10.1101/060012.

97. Hänzelmann, S., Castelo, R. & Guinney, J. GSVA: gene set variation analysis for microarray and RNA-seq data. BMC Bioinformatics 14, 7 (2013).

98. Szolek, A. et al. OptiType: precision HLA typing from next-generation sequencing data. Bioinformatics 30, 3310–3316 (2014).

99. Orenbuch, R. et al. arcasHLA: high-resolution HLA typing from RNAseq. Bioinformatics 36, 33–40 (2020).

100. O’Donnell, T. J., Rubinsteyn, A. & Laserson, U. MHCflurry 2.0: Improved pan-allele prediction of MHC class I-presented peptides by incorporating antigen processing. Cell Syst. 11, 42–48.e7 (2020).

101. Nilsson, J. B. et al. Accurate prediction of HLA class II antigen presentation across all loci using tailored data acquisition and refined machine learning. Sci. Adv. 9, eadj6367 (2023).

102. Altschul, S. F., Gish, W., Miller, W., Myers, E. W. & Lipman, D. J. Basic local alignment search tool. J. Mol. Biol. 215, 403–410 (1990).

## Supplementary References

1. Granados, D. P. et al. Impact of genomic polymorphisms on the repertoire of human MHC class I-associated peptides. Nat. Commun. 5, 3600 (2014).

2. Fuchs, K. J. et al. Expanding the repertoire reveals recurrent, cryptic, and hematopoietic HLA class I minor histocompatibility antigens. Blood 143, 1856–1872 (2024).

3. Smart, A. C. et al. Intron retention is a source of neoepitopes in cancer. Nat. Biotechnol. 36, 1056–1058 (2018).

4. Bykova, N. A., Malko, D. B. & Efimov, G. A. In silico analysis of the minor histocompatibility antigen landscape based on the 1000 genomes project. Front. Immunol. 9, 1819 (2018).

5. Kirby, L. et al. Oligodendrocyte precursor cells present antigen and are cytotoxic targets in inflammatory demyelination. Nat. Commun. 10, 3887 (2019).

6. Aydin, S. et al. Antigen recognition detains CD8+ T cells at the blood-brain barrier and contributes to its breakdown. Nat. Commun. 14, 3106 (2023).

7. Clarkson, B. D., et al. Human neurons stimulated with IFNγ present HLA class I-restricted autoantigens to cytotoxic CD8+ T cells. bioRxivorg (2026) doi:10.64898/2026.05.22.727243.

8. Yang, X. et al. Autoimmunity-associated T cell receptors recognize HLA-B*27-bound peptides. Nature 612, 771–777 (2022).

9. Nelson, R. W. et al. T cell receptor cross-reactivity between similar foreign and self peptides influences naive cell population size and autoimmunity. Immunity 42, 95–107 (2015).

10. Balachandran, V. P. et al. Identification of unique neoantigen qualities in long-term survivors of pancreatic cancer. Nature 551, 512–516 (2017).

11. Ma, Q. et al. Histamine H2 receptor deficit in glutamatergic neurons contributes to the pathogenesis of schizophrenia. Proc. Natl. Acad. Sci. U. S. A. 120, e2207003120 (2023).

12. Shan, L., Fronczek, R., Lammers, G. J. & Swaab, D. F. The tuberomamillary nucleus in neuropsychiatric disorders. Handb. Clin. Neurol. 180, 389–400 (2021).

